# Innate immune memory after brain injury drives inflammatory cardiac dysfunction

**DOI:** 10.1101/2023.10.04.560805

**Authors:** A Simats, S Zhang, D Messerer, J Cao, F Chong, S Besson-Girard, O Carofiglio, S Filser, N Plesnila, C Braun, Ö Gökce, M Dichgans, K Hatakeyama, B Bonev, E Beltrán, C Schulz, A Liesz

## Abstract

The enormous medical burden of stroke is not only due to the brain injury itself and the acute systemic effects, but is largely determined by chronic comorbidities that develop secondarily after stroke. We hypothesized that the high rate of comorbidity developing after a stroke might have a shared immunological cause, however, the chronic effects of brain injury on systemic immunity have so far been barely investigated. Here, we identified myeloid innate immune memory as a cause of remote organ dysfunction after stroke. Using single-cell sequencing, we identified persistent pro-inflammatory transcriptomic changes in resident monocytes/macrophages in multiple organs one month after experimental ischemic brain injury, which was particularly abundant in the heart and associated with the development of cardiac fibrosis and diastolic dysfunction. A similar phenotype was seen in myocardial autopsy samples from stroke versus control patients. We observed chronic functional changes in myeloid hematopoiesis driven by post-stroke IL-1β-mediated epigenetic changes. These alterations could be transplanted to naïve recipient mice and were sufficient to induce cardiac dysfunction. By effectively blocking the trafficking of pro-inflammatory monocytes from the bone marrow to the heart using a dual CCR2/5 inhibitor, we successfully prevented post-stroke cardiac dysfunction. This approach holds promising potential as a novel immune-targeted secondary prevention therapy. We anticipate that the epigenetic immune reprogramming mechanisms detailed here for the brain-heart axis could be generalized to provide a novel framework for explaining the development of various comorbidities after acute tissue injury in remote organs.

## INTRODUCTION

We and others have previously shown that acute brain injuries induce a sterile, systemic inflammatory response.^1^ The inflammatory response to sterile injury is rapidly initiated by the release of immunogenic alarmins, such as nuclear proteins or DNA from necrotic cells to the blood circulation.^2^ It is further characterized by an increase in blood cytokine levels, the mobilization of immune cells and profound changes in immune cell composition and function.^3^ In contrast to the acute inflammatory response within hours to few days, the chronic effects of brain injury on systemic immunity are largely unknown. Few studies mainly investigating blood biomarkers have suggested chronic changes in the concentrations of circulating cytokines and other inflammatory markers, such as Interleukin (IL)-6, IL-1, C-reactive protein (CRP), Interferon (IFN)-γ and High-Mobility Group Box 1 (HMGB1).^4–7^ However, a detailed analysis of the chronically compromised systemic immune compartment after brain injury is still missing and the underlying mechanisms are largely unknown.

Acute brain injuries due to ischemic stroke are a leading cause of mortality and long-term disabilities in adults. Besides the early mortality and morbidity due to the ischemic brain injury itself, long-term morbidity after stroke is also due to the high prevalence of secondary comorbidities and complications, such as cognitive impairment and dementia, post-stroke depression, cardiac events, persistent vascular inflammation and stroke-induced metabolic disturbances.^8–12^ Yet, the exact cause of this increased risk of long-term secondary comorbidities after stroke remains elusive.

Recent studies have demonstrated long-term changes in the function of innate immune cells after bacterial infections or vaccination. This phenomenon has been termed innate immune memory or ‘trained immunity’ in contrast to antigen-specific adaptions in long-lived lymphocytes (T and B cells).^13^ Innate immune memory has been demonstrated in proof-of-concept infection studies to alter the responsiveness to pathogens after re-infection. This represents a beneficial evolutionary mechanism for the clearance of infectious pathogens, but can also result in potentially pathological functions during aging and autoimmunity due to aberrant inflammation.^14^ While the concept of trained immunity has been developed for innate immune memory after infection, barely any information is available for similar mechanisms after sterile tissue injuries. Epigenetic changes in myeloid cells have been reported in models of organ transplantation,^15^ experimental arthritis and in patients with systemic lupus erythematosus.^16,17^ Moreover, alarmin mediators released after sterile tissue injury utilize the same pattern recognition receptors as pathogen-associated molecular patterns, and therefore might instigate similar downstream effects on innate immunity. Therefore, we hypothesized that sterile tissue injuries such as stroke might result in similar long-term innate immune memory as observed after infections, and that these long-term immunological consequences after stroke might drive secondary comorbidities.

Using single-cell sequencing of tissue-resident macrophages and circulating monocytes, we observed substantial transcriptomic changes to a pro-inflammatory phenotype one month after experimental stroke. These changes were present across all peripheral organs. Cardiac monocytes/macrophages showed the most pronounced long-term transcriptomic changes, which were associated with the development of cardiac dysfunction due to fibrosis and increased cardiac inflammation after brain ischemia both in experimental mice and in stroke patients. We identified IL-1β-driven epigenetic changes in the myeloid compartment, which in transplantation experiments was proven to be causative of the long-term cardiac phenotype after stroke. Finally, blocking the recruitment of monocytes after stroke to remote organs prevented secondary cardiomyopathy.

## RESULTS

### Stroke induces long-term inflammatory changes in systemic monocytes/macrophages

To test a potential effect of stroke on long-term systemic inflammation, we performed a comprehensive single-cell mRNA sequencing analysis of CD45+CD11b+ myeloid cells from blood and multiple peripheral organs 1 month after experimental ischemic stroke, which have previously been associated with inflammatory consequences of brain injury in the acute phase (**Fig. 1A**).^2,20–22^ After performing unsupervised clustering, we projected a total of 29,124 myeloid cells and identified 20 independent clusters based on the most variable genes (**Fig. 1B**, Fig. S1A-C). We observed a large number of genes still being differentially regulated at this late chronic time point after stroke particularly within the population of monocytes/macrophages, while other cell populations including neutrophils or dendritic cells were less affected (**Fig. 1C**, Fig. S1D). Transcriptomic changes in monocytes/macrophages after stroke were associated with a pro-inflammatory phenotype, characterized by significant upregulation of various biologically relevant inflammatory signaling pathways of circulating monocytes and tissue macrophages, including increased expression of genes involved in chemotaxis and cell adhesion (e.g. Cx3cr1, Lyz2, Icam1 and Itga4), cytokine-and interferon (IFN)-mediated signaling pathways (e.g. Csf1r, Cebpa, Itgb2, Il10ra, Aim2, Irf8 and Irf5) and pattern recognition receptors (PRRs) (e.g. Irak2 and Tab2) (**Fig. 1D**, Fig. S1E).^18,19^ Furthermore, through a biological network analysis, we confirmed the involvement of other pro-inflammatory mediators, such as interleukin (IL)-12, IL-1 and IFN-α and -β, and the downstream NF-kb and Akt signaling pathways in the activated phenotype of resident monocytes/macrophages in peripheral organs one month after stroke (Fig. S1F).

**Figure 1.**
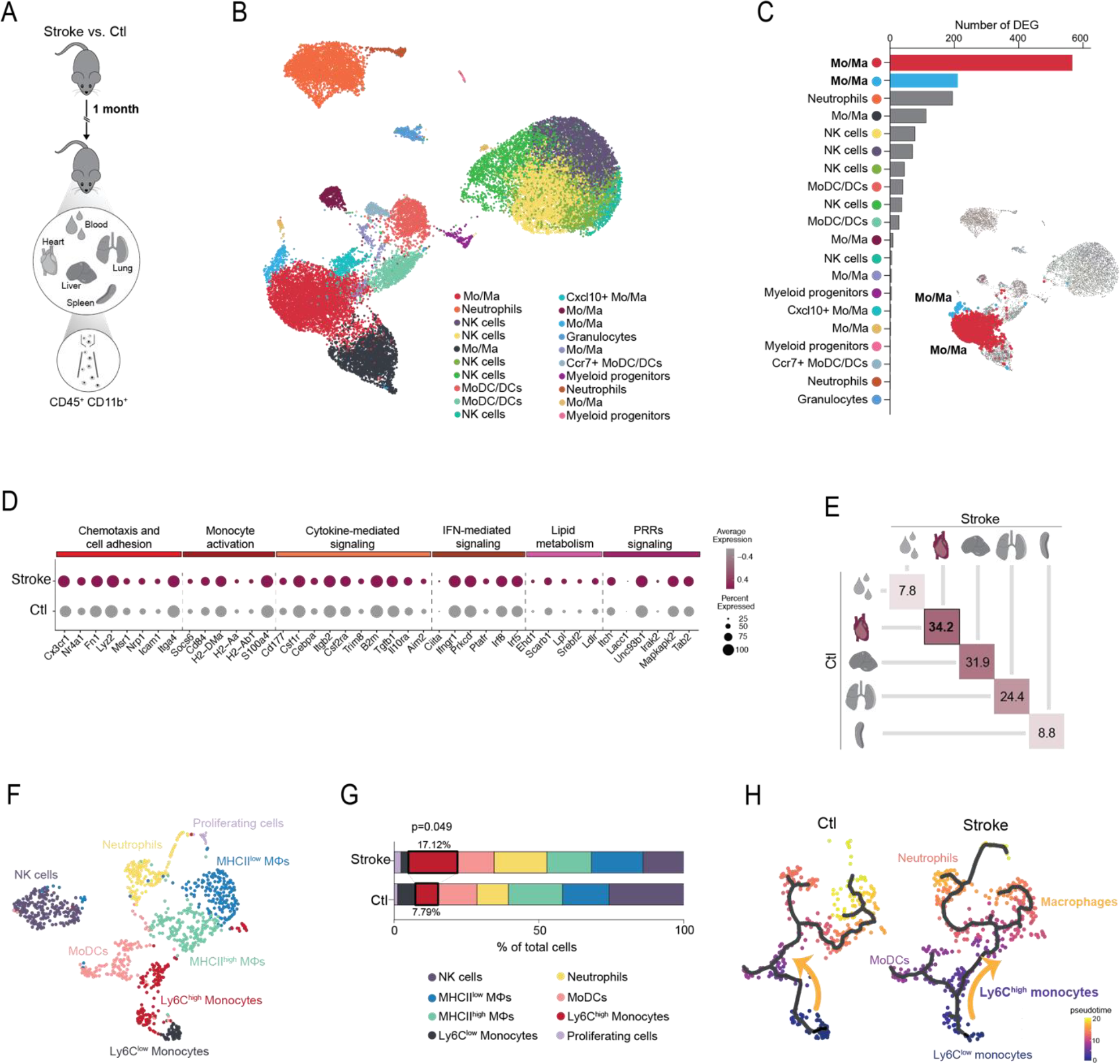
Stroke induces long-term inflammatory changes in systemic monocytes/macrophages. **(A)** Schematic of the experimental design: myeloid cells were sorted from blood and peripheral organs 1 month after experimental ischemic stroke (n=4/group), and the transcriptomic profile of sorted cells was analyzed using single-cell mRNA sequencing. **(B)** Uniform Manifold Approximation and Projection (UMAP) plot of a total of 29,124 CD45^+^ CD11b^+^ cells, colored by identified clusters. **(C)** Number of differentially expressed genes (DEG) between stroke and control conditions per identified population (adjusted p value <0.05). Two clusters of monocytes/macrophages show the highest number of DEG between conditions (565 and 212, respectively). **(D)** Dot plot showing the expression levels of selected genes in the 2 clusters of monocytes/macrophages highlighted in (C) between conditions. Genes were selected from the enriched GO terms in the set of DEG between stroke and control mice in the 2 selected populations of monocytes/macrophages (adjusted p value <0.1). The dot size corresponds to the fraction of cells within each condition expressing the indicated transcript, and the color indicates average expression. **(E)** Euclidian distances in the PCA space between the stroke and control samples per each individual organ. The PCA was calculated from a total of 18,834 genes identified in all CD45^+^CD11b^+^ cells. Distances were calculated from the mean point of each cluster (group), composed of four individual samples each. **(F)** UMAP plot of 1,117 CD45^+^ CD11b^+^ cells from the heart, colored by identified populations. **(G)** Stacked bar graph showing the percentage of cells of each identified population per condition (chi-square test). **(H)** Monocle3 pseudo-temporal ordering of CD45^+^ CD11b^+^ cells from the heart superimposed on the UMAP plot and split by condition. Cells are colored based on their progression along pseudo-temporal space.

Across organs, changes in the monocyte/macrophage population where particularly pronounced in the heart and liver. Principal component analysis based on a total of 18.835 genes showed that changes in blood, spleen, and lung were more subtle (**Fig. 1E**, Fig. S1G). Specifically, we detected the selective expansion of a Ly6C^high^-expressing monocyte population in hearts 1 month after stroke (**Fig. 1 F,G**, Fig. S1H), which changed the differentiation trajectories of monocytes to cardiac macrophages (**Fig. 1H**). In addition, cardiac Ly6C^high^ monocytes after stroke showed increased expression of genes related to tissue residency when compared to circulating monocytes (Fig. S1I). Taken together, these results suggest that stroke chronically promotes the recruitment of circulating Ly6C^high^ monocytes to the heart, which might further differentiate into tissue-resident macrophages.

### Stroke results in chronic cardiac diastolic dysfunction

To assess the functional consequences of changes in the cardiac monocyte/macrophage population during the chronic phase following stroke, we investigated cardiac function using Doppler echocardiography. We observed a persistent reduction in end-diastolic left ventricle (LV) volume, while the systolic function, measured by the ejection fraction and fractional shortening, was only transiently affected by stroke in the acute phase (**Fig. 2A**, Fig. S2A). These findings suggest the selective development of chronic diastolic dysfunction, which was further confirmed using pulse wave doppler of the apical four-chamber-window demonstrating compromised left ventricular compliance by decreased mitral valve E wave deceleration time (Fig. S2B). Heart failure in patients with preserved ejection fraction but diastolic dysfunction is commonly associated with cardiac fibrosis, which impairs the rapid LV filling due to increased myocardial stiffness.^23,24^ Hence, we evaluated the composition of the extracellular matrix (ECM) in hearts one month after stroke or in control mice and observed significantly increased LV fibrosis mainly due to increased deposition of Type I collagen (**Fig. 2B,C**, Fig. S2C,D). Further analysis of collagen orientation in the LV myocardium by second harmonic generation microscopy revealed increased fiber disorganization after stroke, suggesting ECM remodeling in addition to increased deposition (**Fig. 2D**). Therefore, we analyzed cardiac matrix metalloproteinase (MMP) activity—key effector enzymes in ECM remodeling—in stroke and control mice and observed significantly increased MMP9 activity by gel zymography (**Fig. 2E**) as well as total pro-MMP9 protein content (Fig. S2E). More specifically, we detected by smFISH significantly increased *Mmp9* transcripts in cardiac macrophages, as the most likely source of increased cardiac MMP9 expression, which was confirmed by rt-PCR of sorted cardiac monocytes/macrophages (**Fig. 2F**, Fig. S2F). Corresponding to results of the single-cell sequencing analysis (see Fig. 1F,G), we also observed by flow cytometry an increased number of CCR2^+^ cardiac macrophages in post-stroke hearts (**Fig. 2G**, Fig. S2G), suggesting a higher infiltration of circulating monocytes and an enhanced monocyte-to-macrophage differentiation chronically after stroke.^25,26^ In addition, we observed that circulating monocytes after stroke also have a significantly increased expression of *Mmp9* (**Fig. 2H**), which suggests a causative role of infiltrating monocytes on the enhanced MMP9 activity in the post-stroke heart.

**Figure 2.**
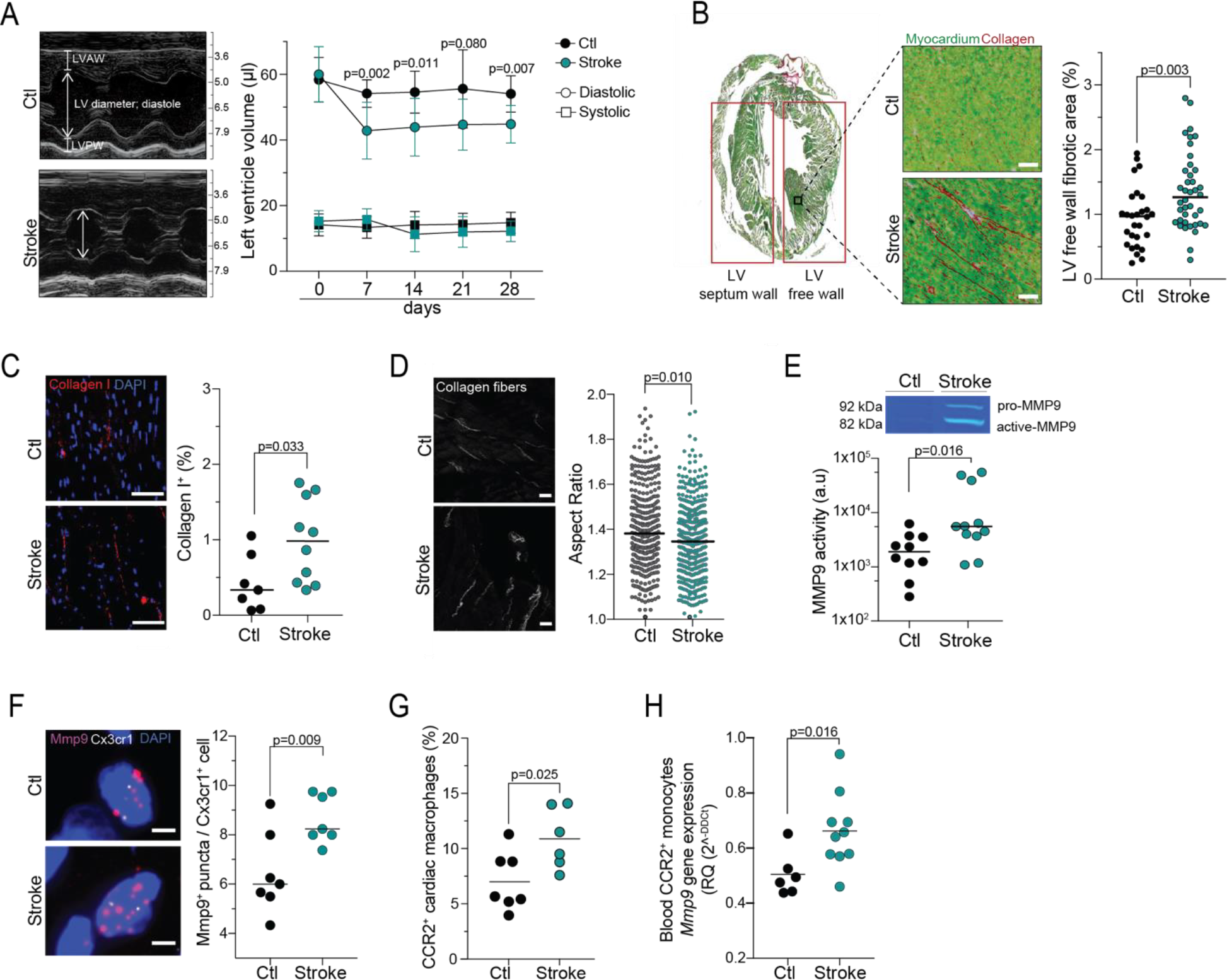
Stroke results in chronic cardiac diastolic dysfunction and inflammatory cardiac remodeling. **(A)** Representative ultrasound images (M-mode) performed at day 0, 7, 14, 21 and 28 after stroke (n = 12) and control mice (n = 6). Corresponding quantification of the left ventricle (LV) volume in systole (squares) and diastole (circles). LVAW: Left ventricle anterior wall; LVPW: left ventricle posterior wall. **(B)** Representative images of the Sirius red/Fast green collagen staining performed on cardiac coronal sections for stroke and control mice. Corresponding quantification of cardiac fibrosis in the LV free wall (t test, n=7/10 per group, 4 independent sections per mouse). **(C)** Representative image of immunofluorescence staining for the detection of collagen I in heart coronal sections (left; scale bar = 50 µm). Corresponding quantification of collagen I content, expressed in percentage of total area of the LV free wall (right; t test, n = 8/10 per group). **(D)** Representative second harmonic generation (SHG) images for the detection of the organization of fibrillar collagen in the heart (left; scale bar = 20 µm). Corresponding quantification of the Aspect ratio (right; nested t test, n = 9/10 mice per group, 35-50 images per mouse heart). **(E)** Enzymatic MMP9 activity was measured by *in gel* zymography of heart samples from stroke and control mice (U test, n = 10/11 per group). **(F)** Representative images of single molecule fluorescence *in situ* hybridization (smFISH) for the detection of *Mmp9* mRNA expression in *Cx3cr1*^+^ cardiac cells from stroke and control mice (left, scale bar = 5 µm). Corresponding quantification of the number of *Mmp9* mRNA puncta per *Cx3cr1*^+^ cell. DAPI was used as nuclear dye (right; t test, n = 7 per group). **(G)** Cardiac myeloid cells were isolated from the heart of stroke and control mice 1 month after stroke and analyzed by flow cytometry. Quantification of total counts for CCR2^+^F4/80^+^ cells in the heart of stroke and control mice (t test; n=6/8 per group). **(H)** RT-qPCR was performed on sorted blood CCR2+ monocytes to measure the expression levels of *Mmp9* mRNA, quantified relative to the expression of the housekeeping gene encoding *Ppia* and normalized to control levels (U test, n=6/10 per group).

We next aimed to validate the translational relevance of these observations. For this, we obtained myocardial autopsy samples from patients that had died 1-3 months after stroke or from age-matched control subjects that had died without cardiac or brain disorder (confirmed by autopsy, **Fig. 3A**). We observed significantly increased ECM deposition in the left ventricular wall of stroke patients (**Fig. 3B,C**) and detected a significant increase in CCR2^+^ monocyte counts in post-stroke hearts, which correlated with cardiac collagen content, while total monocyte and macrophage counts did not differ between groups (**Fig. 3D-F**). Consistent with our experimental results, human cardiac macrophages also expressed significantly more *MMP9* transcripts after stroke (**Fig. 3G**). Finally, we performed bulk mRNA sequencing on adjacent tissue sections used for the histological analyses above and found significant transcriptional differences between control subjects and after stroke (**Fig. 3H,I**). Interestingly, several of the upregulated genes in stroke patients were associated with extracellular matrix remodeling including *PON1* (paraoxonase 1) and *KRTCAP2* (Keratinocyte-associated protein 2). Taken together, we observed marked cardiac fibrosis and ECM remodeling in both experimental mice and stroke patients, which was further associated with diastolic dysfunction after experimental stroke. A hallmark of this secondary cardiac pathology after stroke is the increased recruitment and proinflammatory profile of cardiac monocytes/macrophages.

**Figure 3.**
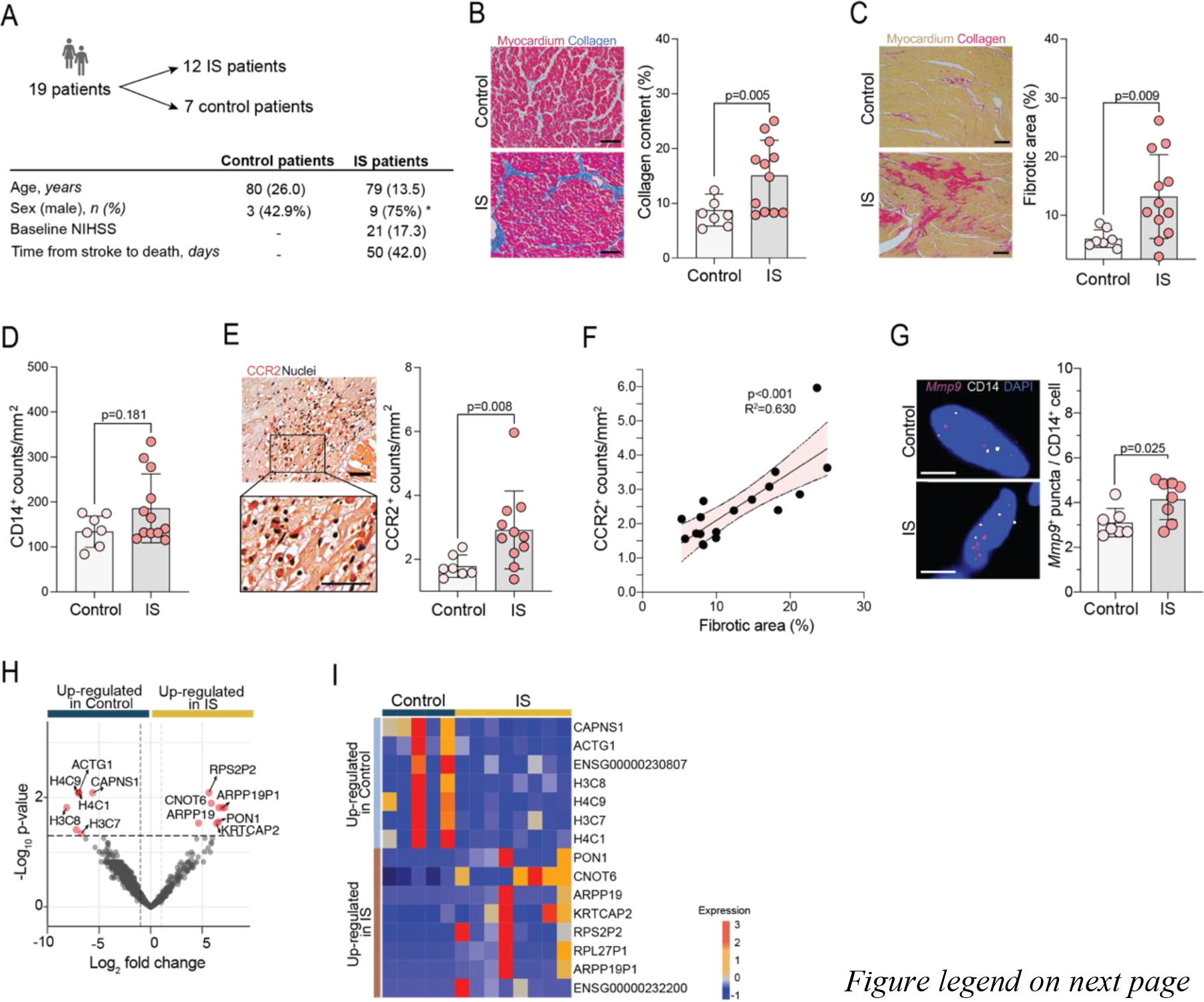
Stroke increases cardiac fibrosis and monocyte accumulation in patients. **(A)** Myocardial autopsy samples were collected from a total of 19 patients: 12 ischemic stroke (IS) patients and 7 controls, who died without cardiac or brain disorder (confirmed by autopsy). Basic demographical and clinical characteristics are depicted. Data is expressed as median (IQR), unless stated otherwise. *p<0.05 (chi-square test). **(B)** Representative images of the Masson Trichrome staining (left, scale bar = 0.1 mm). Corresponding quantification of the collagen content, assessed with the Masson Trichrome staining, expressed as percentage of total cardiac area (mid, t test). **(C)** Representative images of the Sirius red/Fast green collagen staining (left, scale bar = 0.2 mm). Corresponding quantification of the collagen content, assessed with the Sirius red/Fast green collagen staining, expressed as percentage of total cardiac area (right, t test). **(D)** Quantification of CD14^+^ cells (t test). **(E)** Representative image (left, scale bar = 0.05 mm) and quantification (right, t test) of CCR2^+^ cells by immunohistochemistry. **(F)** Correlation of CCR2^+^ cell counts with cardiac fibrosis, assessed with the Masson Trichrome staining (Pearson correlation test). **(G)** Representative images of single molecule fluorescence *in situ* hybridization (smFISH) for the detection of *Mmp9* mRNA puncta expression in *CD14*^+^ cardiac cells from stroke and control patients (left, scale bar = 5 µm). Corresponding quantification of the number of Mmp9 mRNA puncta per CD14+ cell. DAPI was used as nuclear dye (right; t test, n = 7/9 per group). **(H)** Volcano plot showing regulated genes in the myocardium between IS and control patients. Colored genes are p<0.05. **(I)** Heatmap showing the TOP differentially expressed genes in the heart between IS and control patients.

### Bone marrow cellularity and function is chronically altered after stroke

To explore the potential mechanisms of post-stroke immune mediated cardiac dysfunction, we performed an in-depth analysis of the myeloid compartment in the chronic phase after stroke. This was motivated by the finding that, based on in vivo EdU-labeling experiments, more than 95% of cardiac myeloid cells will have been replaced within one month after stroke (**Fig. 4A**). Indeed, using an inducible myeloid progenitor reporter mouse strain (Ms4a3^creERT2^xAi14; Fig. S3A-C), we observed substantial cellular recruitment of cardiac monocytes/macrophages with turn-over of approx. 20% of cardiac monocytes/macrophages within 2 weeks (**Fig. 4B**). Therefore, we analyzed long-term effects of stroke on the bone marrow by single cell sequencing, identifying 21 cell clusters and the respective differentiation trajectories from hematopoietic stem and progenitor cells (HSPCs) to all other mature myeloid populations (**Fig. 4C,D**, Fig. S3D,E). Subsequently, we computed the number of differentially expressed genes (DEGs) between stroke and control conditions for each of these populations, revealing a distinct transcriptomic signature in Ly6C^high^ monocytes chronically after stroke (**Fig. 4E**). Additionally, we observed a substantial number of DEGs in HSPCs, suggesting that not only mature monocytes but also progenitor populations exhibited persistent transcriptional alterations during the chronic stage after stroke. To validate these findings, flow cytometry analyses were conducted, confirming altered bone marrow cellularity characterized by increased numbers of mature monocytes and neutrophils, as well as significantly increased myeloid progenitor populations after stroke (**Fig. 4F,G**, Fig. S3F). Labeling of proliferating cells with a single intraperitoneal EdU injection one month after stroke indicated significantly increased proliferation rates of myeloid progenitor cell populations as the most likely cause of the chronically altered BM cellularity after stroke (**Fig. 4H**).

**Figure 4.**
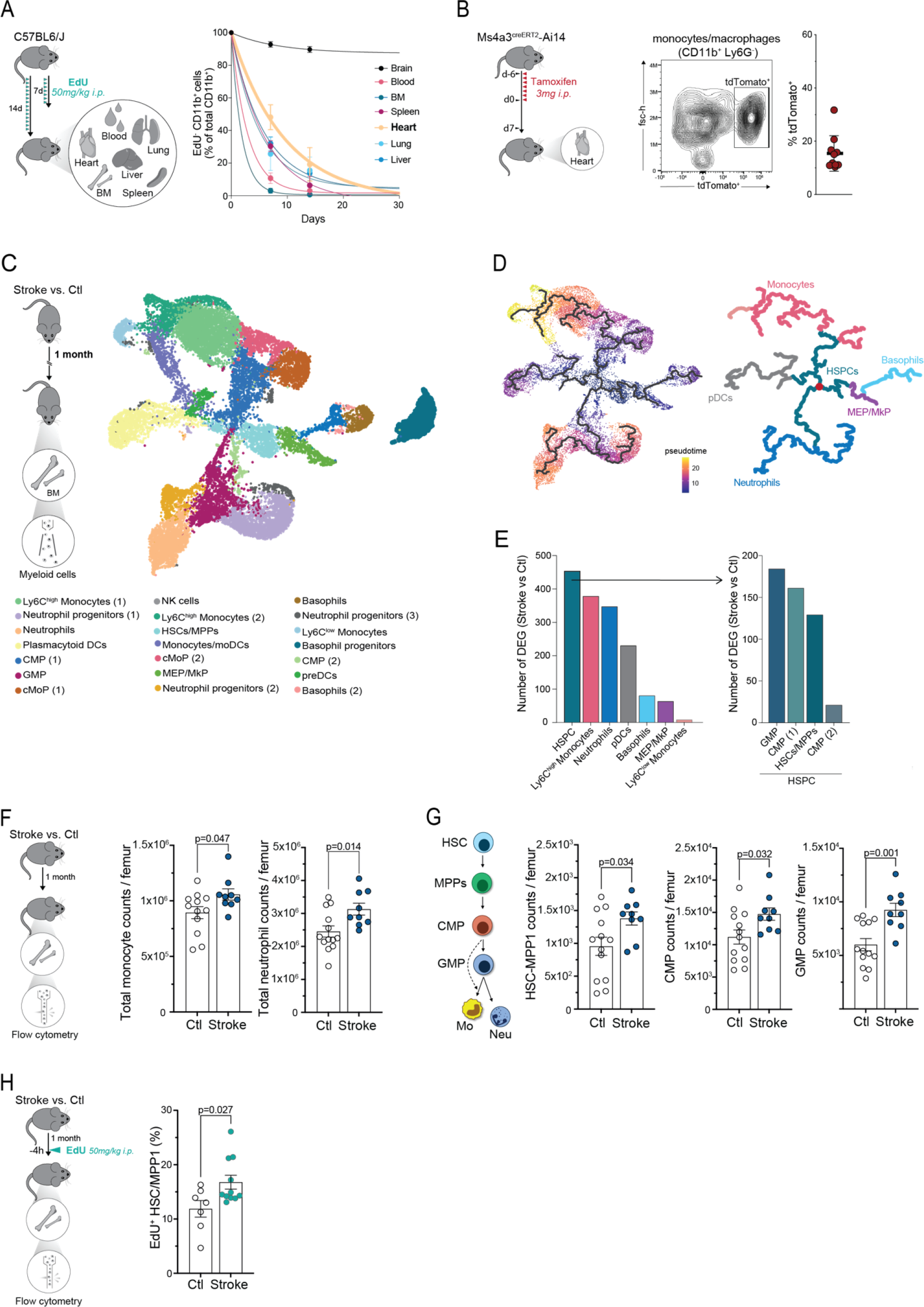
Bone marrow cellularity and function are chronically altered after stroke. **(A)** Schematic experimental design: stroke mice were daily administered with EdU for 7 or 14 days after stroke. Lineage negative myeloid cells from the blood, and peripheral organs were analyzed by flow cytometry (left). Corresponding quantification of the percentage of EdU^+^ CD11b^+^ cells over time (right) (n=6/7 per organ). **(B)** Ms4a3^creERT2^-Ai14 mouse were treated with tamoxifen for 7 consecutive days. Mice were sacrificed 1 week later and the myocardial myeloid cells were analyzed by flow cytometry (left panel). Representative gating strategy for the TdT^+^ cardiac monocytes/macrophages (CD45^+^ Ly6G^-^ CD11b^+^, mid panel) and quantification of the percentage of TdT^+^ cardiac monocytes/macrophages (right panel, n=5). (**C**) Schematic experimental design: lymphoid lineage-(CD3, CD4, CD8a, CD19 and Ter119) and neutrophil-(Ly6G) negative myeloid cells were sorted from the BM of control and stroke mice one month after stroke (n=8). The transcriptomic profile of sorted lineage-negative cells was analyzed using single-cell mRNA sequencing (left panel). UMAP plot of the 22,169 myeloid cells sorted from the BM of control and stroke mice (right panel). **(D)** Monocle3 pseudo-temporal ordering of myeloid BM cells, superimposed on the UMAP plot. Cells are colored based on their progression along pseudo-temporal space (left panel) or by cell types (right panel). **(E)** Number of differentially expressed genes (DEG) between stroke and control conditions per cell type (adjusted p value <0.05). **(F)** Schematic experimental design: BM cells were isolated from stroke and control mice 1 month after stroke and analyzed by flow cytometry (left panel). Corresponding monocyte and neutrophil cell counts were quantified in stroke and control mice (mid and right panels) (U-test; n=9/13 per group). **(G)** Schematic of the differentiation path of hematopoietic stem cells (HSC) towards monocytes (Mo) and neutrophils (Neu) (left panel). Corresponding cell count quantifications of HSC-multipotent progenitors 1 (MPP1), common myeloid progenitors (CMP) and granulocyte-monocyte progenitors (GMP) in stroke and control mice (mid and right panels) (U-test; n=9/13 per group). **(H)** Schematic experimental design: stroke and control mice were administered with EdU 4h before sacrifice. Subsequently, BM cells were isolated and analyzed by flow cytometry (left panel). Corresponding quantification of the percentage of EdU^+^ cells from HSC/MPP1 in stroke and control mice (U-test; n=7/11 per group).

### Stroke induces persistent innate immune memory

To test the persistence and potentially causal role of the observed myeloid changes chronically after stroke, we transplanted enriched progenitor and stem cells from bone marrows one month after stroke or control mice into naïve recipients, using a genetic BM depletion model (poly(I:C) administration to Mx1^Cre^:c-myb^fl/fl^ mice) in order to avoid confounding effects by irradiation or chemotherapy (**Fig. 5A**). One month after transplantation, we isolated eGFP-positive transplanted myeloid cells for single-cell mRNA sequencing and confirmed successful BM repopulation for all analyzed animals, without differences in the BM repopulation potential between groups (**Fig. 5B**, Fig. S4A-F). Differential gene expression between stroke and control transplanted BM cells showed substantial transcriptomic differences one month after transplantation in the naïve recipient mice (Fig. S4G,H). Interestingly, comparing the most significantly regulated transcriptomic pathways between the transplanted groups to the original difference one month after stroke and control revealed a highly conserved phenotype in the transplanted cells after stroke (**Fig. 5C**). Both BM cells from stroke mice and BM cells from recipient mice transplanted with stroke BM cells retained a pro-inflammatory activated phenotype and showed sustained high expression of genes involved in chemotaxis and cell adhesion, monocyte activation, and cytokine- and interferon (IFN)-mediated signaling pathways. Projecting all individual samples into a PCA bidimensional space, we found that cells from mice transplanted with stroke BM showed spatial proximity to cells from stroke mice (Fig. S4I), confirming that myeloid cells acquire a distinct pro-inflammatory phenotype after stroke that is transmissible by BM transplantation.

**Figure 5.**
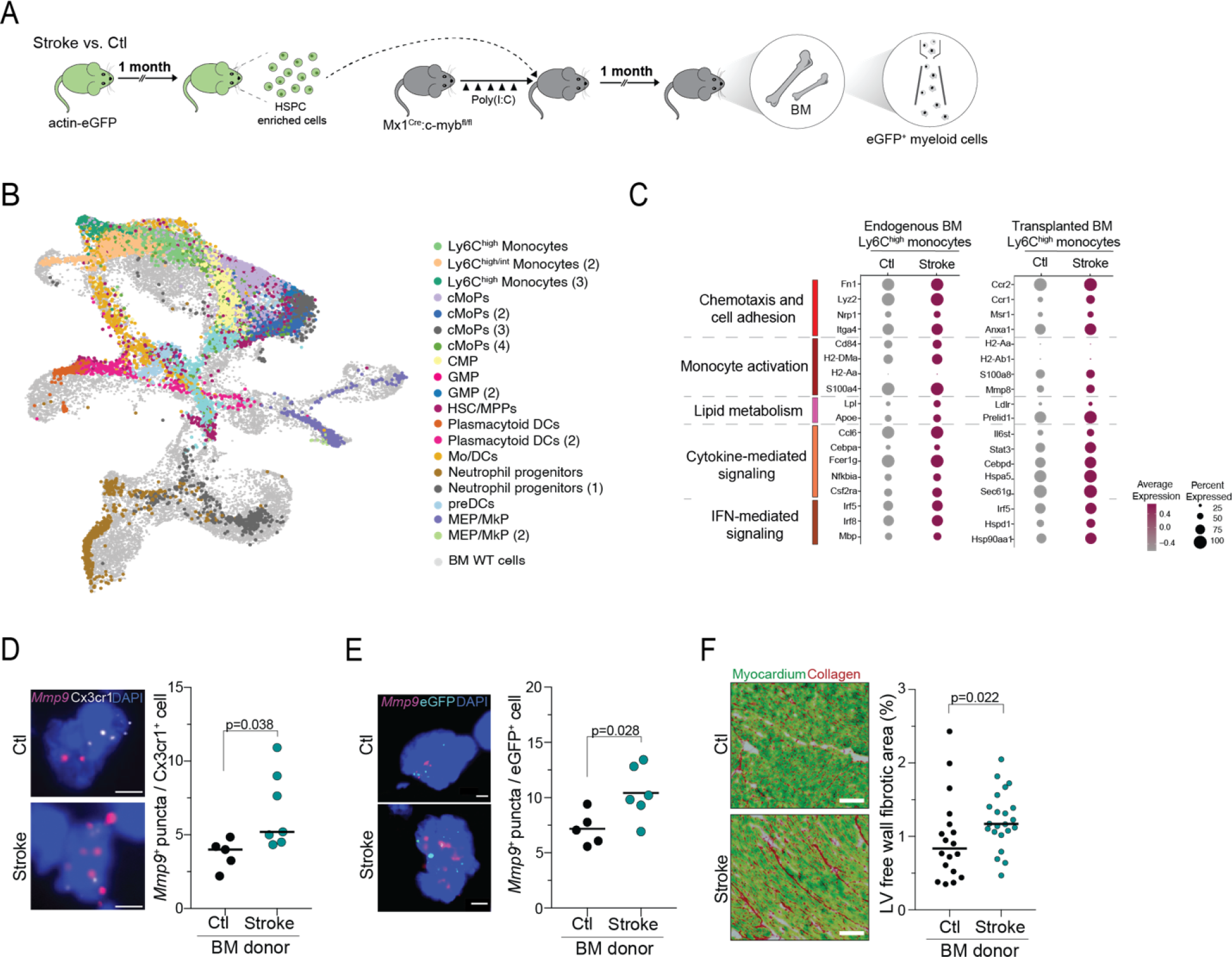
Stroke induces stable innate immune memory. **(A)** Schematic experimental design: bone marrow (BM) cells enriched for HSPCs were isolated from stroke and control actin-eGFP mice and transplanted into BM-depleted Mx1^Cre^:c-myb^fl/fl^ mice. One month after transplantation, mice were sacrificed and eGFP-positive myeloid cells were isolated and analyzed using single-cell mRNA sequencing. **(B)** UMAP plot of 25,358 myeloid cells from the BM of transplanted mice, colored by identified populations and superimposed on the UMAP plot of the myeloid cells from the endogenous BM from control and stroke mice (cells in gray, Fig. 4C for reference). **(C)** Dot plot showing the expression of selected genes in the Ly6C^high^ monocytic populations from the endogenous BM of stroke and control mice (left column) and the BM of recipient mice transplanted with stroke and control eGFP^+^ HSPC-enriched BM cells (right column). Genes were selected from the enriched GO terms in the set of DEG between stroke and control mice in the populations of Ly6C^high^ monocytes (adjusted p value<0.1). The dot size corresponds to the fraction of cells within each condition expressing the indicated transcript, and the color indicates average expression. **(D)** Representative images of single molecule fluorescence *in situ* hybridization (smFISH) for the detection of *Mmp9* mRNA expression in myeloid cells from the heart of recipient mice transplanted with stroke and control BM cells (left, scale bar = 5 µm). Corresponding quantification of the number of *Mmp9* puncta per *Cx3cr1*^+^ cardiac cell. DAPI was used as nuclear dye (U test, n=5/7 per group). (**E**) Representative images of single molecule fluorescence *in situ* hybridization (smFISH) for the detection of *Mmp9* mRNA expression in eGFP^+^ cardiac cells from recipient mice transplanted with stroke and control BM cells (left, scale bar = 5 µm). Corresponding quantification of the number of *Mmp9* puncta per eGFP^+^ cell. DAPI was used as nuclear dye (U test, n=5/6 per group). **(F)** Representative images of the Sirius red/Fast green collagen staining performed on cardiac coronal sections from recipient mice transplanted with stroke and control BM cells. Quantification of cardiac fibrosis in the left ventricle (LV) free wall (t test, n=5/7 per group, 3/4 heart sections per mouse).

Next, we analyzed the hearts of BM recipient mice for hallmarks of ECM remodeling observed after stroke. We found significantly increased *Mmp9* expression in cardiac macrophages of mice receiving BM from stroke donors in comparison to control donors and more specifically, an increase of *Mmp9* expression particularly in transplanted GFP-positive cardiac monocytes/macrophages (**Fig. 5 D,E**). Moreover, we observed significantly increased cardiac fibrosis in animals receiving the stroke BM transplant in comparison to control BM recipients (**Fig. 5F**). Together, these findings demonstrate that myeloid function is not only stably altered after stroke but that these changes in the myeloid compartment are sufficient to drive secondary cardiac fibrosis.

### Innate immune memory is mediated by early post-stroke Interleukin-1β secretion

We have previously identified systemic inflammasome activation and subsequent secretion of Interleukin (IL)-1β as a hallmark of the acute inflammatory response to tissue injury including stroke, with serum IL-1β concentrations peaking within the first hours after injury.^2^ Studies in experimental infection models have suggested IL-1β-mediated effects to be involved in epigenetic reprogramming and innate immune memory following infections.^17^ Therefore, we tested the effect of neutralizing the early secretion of IL-1β in response to stroke by systemic administration of IL-1β-specific monoclonal antibodies on bone marrow composition and transcriptomic features one month later. By flow cytometry, we observed efficient normalization of bone marrow cellularity to levels of control mice in animals receiving IL-1β neutralization (**Fig. 6A**, Fig. S5A,B). To further test the hypothesis of potential epigenetic changes in myeloid cells by the acute IL-1β secretion after stroke, we performed single-cell ATAC sequencing for analysis of open chromatin accessibility in control mice and in animals one month after stroke which either received IL-1β neutralizing antibodies or vehicle control 1h before and after stroke induction. We obtained a total of 13,520 nuclei from Lin^-^ CD45^+^CD11b^+^ BM cells, and identified a total of 11 clusters which were superimposed on the BM mRNA sequencing UMAP plot (Fig. 4B), confirming coverage of the complete BM cell heterogeneity (**Fig. 6B**, Fig. S5C-E). We identified in the HSPC as well as in the mature Ly6C^high^ monocyte clusters distinct differences in chromatin accessibility one month after stroke in comparison to control mice (**Fig. 6C**). Next, we particularly focused on potentially important cell-type-specific differentially-active regulatory sequences,^27^ and identified significant changes in DNA transcription factor (TF) motifs between stroke and control conditions (**Fig. 6D**). This analysis revealed that stroke induced a pronounced alteration in the activity of several TF in HSPC, including CTCF, ETV4 and RUNX2, which have been previously described to regulate HSPC function.^28–30^ Similarly, stroke also changed the epigenetic landscape of mature Ly6C^high^ monocytes, by changing the motif accessibility associated with TF involved in stress and immune responses, such as CTCF, NRF1 (also known as NFE2) and FOS:JUND(AP1).^31–36^ Importantly, we observed that post-stroke IL-1β neutralization prevented most of the stroke-induced changes in chromatin accessibility and attenuated post-stroke differential TF motif activities in both, HSPC and mature Ly6C^high^ monocytes. To explore the functional relevance of the stroke-regulated TFs, we identified genes differentially expressed between control and stroke conditions that were significantly associated with DNA sequences regulated by these transcription factors (**Fig. 6E**). These genes were associated with a number of functional pathways in inflammation and cytokine signaling (**Fig. 6F**). Finally, we observed that neutralization of the early IL-1β release was sufficient to prevent the long-term cardiac phenotype by significantly reducing *Mmp9* expression of cardiac monocytes/macrophages and cardiac fibrosis to levels of control mice without a stroke (**Fig. 6G,H**). These findings confirm the hypothesis that IL-1β-driven epigenetic changes leading to innate immune memory can be causally involved in mediating remote organ dysfunction after stroke.

**Figure 6.**
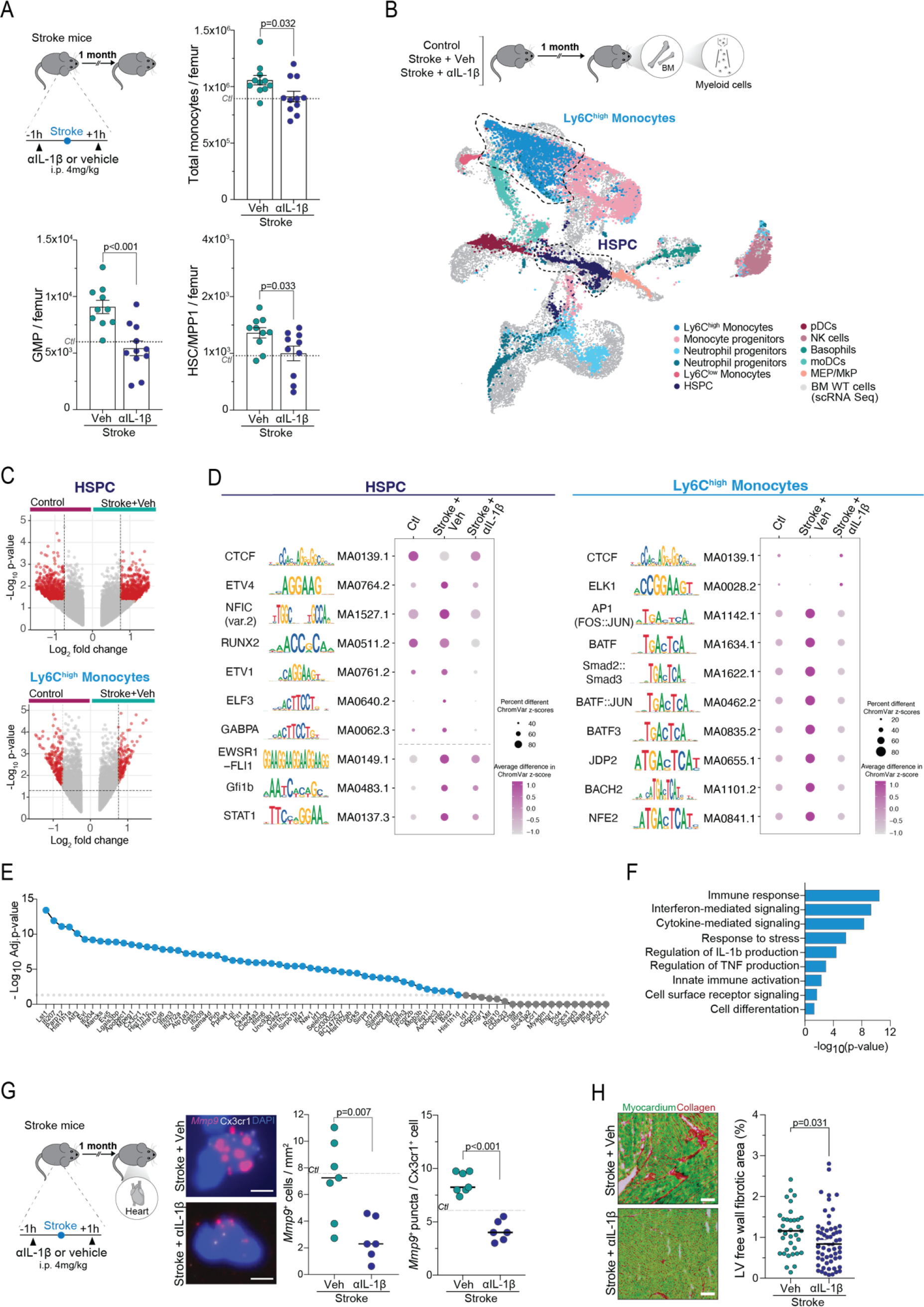
Innate immune memory after stroke is mediated by IL-1β. **(A)** Schematic experimental design: mice received Interleukin (IL)-1β neutralizing antibodies or vehicle 1h before and 1h after stroke induction. One month later, bone marrows (BM) were collected for flow cytometry. Corresponding cell count quantifications of total monocytes, granulocyte-monocyte progenitors (GMP) and HSC-multipotent progenitors 1 (HSC-MPP1) in BM from stroke mice treated with anti-IL-1β (stroke+αIL-1β) or vehicle (stroke+Veh) (U-test; n=10/11 per group). **(B)** Schematic experimental design: nuclei were isolated from lineage-negative myeloid cells sorted from the BM of control and anti-IL1β or vehicle-treated stroke mice 1 month after stroke (n=3/group). The transcriptomic profiles of sorted nuclei were analyzed using single-nuclei ATAC-sequencing (upper panel). UMAP plot of 13,520 myeloid nuclei, colored by identified populations and superimposed on the UMAP plot of the myeloid cells from the endogenous BM of control and stroke mice (cells in gray, Fig. 4C). (**C**) Volcano plots showing the differentially accessible peaks between vehicle-treated stroke and control conditions for HSPC (top panel) and Ly6C^high^ monocytes (bottom panel). Colored peaks are p<0.05 and log2FC > 0.75. **(D)** Dot plot showing per-cell differential motif activity scores between experimental conditions in HSPC (left panel) and Ly6C^high^ monocytes (right panel). Identified motifs with the highest and lowest activity scores between control and vehicle-treated strokes are represented (adjusted p value <0.05). **(E)** Dotplot showing the statistical significance of the differential expression of a list of genes with a significant correlation with the DNA sequences from the top 10 transcription factors with highest differential activity between stroke and control conditions in Ly6C^high^ monocytes. (**F**) Pathway analysis performed by g:Profiler using the differential expressed genes between stroke and control conditions that showed significant correlation with the top 10 transcription factors with highest differential activity between these two conditions in Ly6C^high^ monocytes. Biological processes were grouped and sorted by p value. **(G)** Schematic experimental design: stroke mice received IL-1β neutralizing antibodies or vehicle 1h before and 1h after stroke induction. One month later, hearts were collected for histology (left panel). Representative images of single molecule fluorescence *in situ* hybridization (smFISH) for the detection of *Mmp9* mRNA expression in *Cx3cr1*^+^ cardiac myeloid cells from anti-IL1β and vehicle-treated stroke mice (scale bar = 5 µm). Corresponding quantification of the total *Mmp9*^+^ cardiac cells and the number of *Mmp9* puncta per *Cx3cr1*^+^ cell . DAPI was used as nuclear dye (U test, n=6/7 per group). **(H)** Representative images of the Sirius red/Fast green collagen staining performed on cardiac coronal sections from anti-IL1β and vehicle-treated stroke mice (left panel). Quantification of cardiac fibrosis in the left ventricle (LV) free wall in anti-IL1β or vehicle-treated stroke mice (right panel, t test, n=8/14 per group, 4 heart sections per mouse).

### Blocking BM-to-heart trafficking of monocytes prevents post-stroke cardiac dysfunction

Despite its striking efficacy on innate immune memory and cardiac secondary comorbidities, the translational use of neutralizing IL-1β in stroke patients is limited by an increased risk of infections as revealed by recent clinical trials using IL-1β-neutralizing strategies.^37^ Therefore, we sought for alternative options that could limit secondary remote organ pathologies due to innate immune memory. To block the migration of pro-inflammatory reprogrammed myeloid cells to secondary organs such as the heart after stroke we tested the dual C-C chemokine receptors type 2 and 5 antagonist Cenicriviroc (CVC), which has been developed and proven safe for HIV infections and for steatohepatitis.^38–40^ We found that daily treatment with CVC after experimental stroke substantially reduced monocyte recruitment to the heart (**Fig. 7A,B**, Fig. S6A,B). This effect is likely attributable to blocking the chemokine-dependent invasion of monocytes to secondary organs because CVC increased circulating Ly-6C^high^ monocyte counts and had no direct cytotoxic effect on monocytes (Fig. S6C,D). Consequently, CVC treatment significantly reduced *Mmp9* expression by cardiac monocytes/macrophages and reduced cardiac fibrosis (**Fig. 7C,D** Fig. S6E). Importantly, this therapeutic reduction in inflammatory cardiac ECM remodeling by CVC also significantly improved diastolic cardiac function in the chronic phase after stroke to comparable levels of control mice without a stroke (**Fig. 7E**). In summary, these results demonstrate the therapeutic potential of blocking the BM-to-heart migration of pro-inflammatory programmed monocytes as a means to prevent secondary cardiac comorbidity and alleviate inflammatory cardiac fibrosis following stroke.

**Figure 7.**
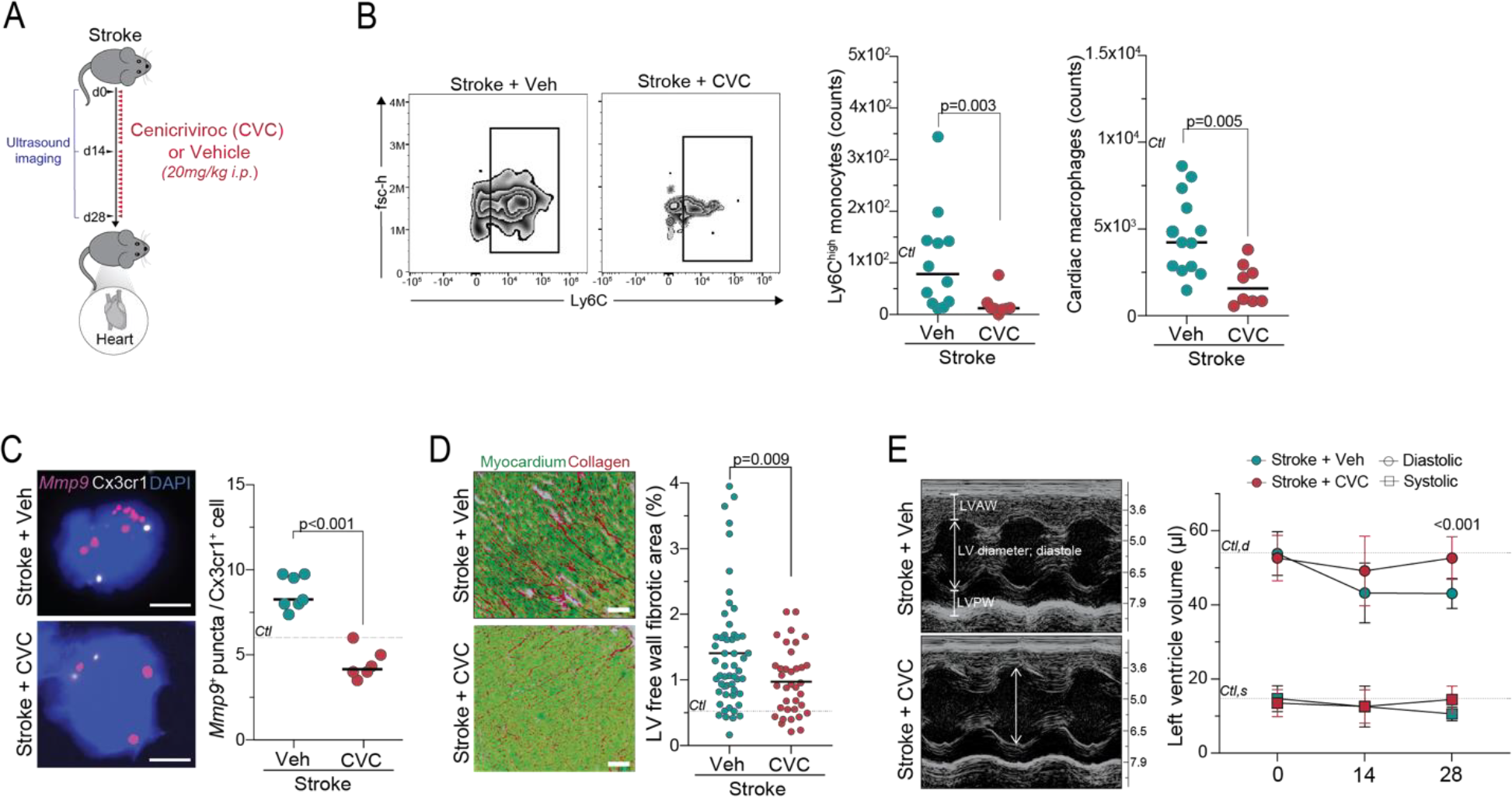
Blocking BM-to-heart migration of monocytes prevents post-stroke cardiac dysfunction. **(A)** Schematic experimental design: Stroke mice were administered daily with the dual C-C chemokine receptors type 2 and 5 antagonist Cenicriviroc (Stroke+CVC) or vehicle (Stroke+Veh) for 28 days. Cardiac ultrasound imaging was performed at day 0, 14 and 28. Mice were sacrificed at day 28 and the hearts were collected for flow cytometry and histological analysis. **(B)** Representative image of the gating for Ly6C^high^ monocytes in CVC- and vehicle-treated stroke mice (left panel) in flow cytometry and corresponding quantification (right panel, U-test; n=8-12 per group). **(C)** Representative smFISH images for the detection of *Mmp9* mRNA expression in cardiac myeloid cells from CVC- and vehicle-treated stroke mice (left panel). Corresponding quantification of the number of *Mmp9* puncta per *Cx3cr1*^+^ cardiac myeloid cell (right panel). DAPI was used as nuclear dye (U test, n=6/7 per group). **(D)** Representative images of the Sirius red/Fast green collagen staining performed on cardiac coronal sections from CVC- and vehicle-treated stroke mice (left panel). Quantification of cardiac fibrosis in the LV free wall in CVC- and vehicle-treated stroke mice. Dashed line indicates mean percentage of fibrotic area in control mice (t test, n=9/14 per group, 4 heart sections per mouse). **(E)** Representative ultrasound images (M-mode) performed at day 0, 14 and 28 after stroke on CVC- and vehicle-treated stroke mice (left panel). Quantification of the left ventricle (LV) volume in systole (squares) and diastole (circles) in CVC- and vehicle-treated stroke mice (right panel). Dashed line indicates mean LV volume in control mice at day 28. Multiple t tests, n=6/12 per group. LVAW: Left ventricle anterior wall; LVPW: left ventricle posterior wall.

## DISCUSSION

Systemic inflammation following a stroke has been identified as a critical factor affecting the short- and long-term prognosis of stroke patients.^41,42^ Interestingly, many of the pre-existing or acquired health conditions that arise after a stroke share common inflammatory mechanisms. These mechanisms can potentially exacerbate the development of other medical complications, leading to a worsened long-term outcome. Consequently, addressing systemic inflammation has emerged as a novel focus for translational research, with initial clinical trials already conducted to minimize functional disabilities in patients and prevent secondary complications.^3^ Our study reveals that stroke triggers persistent inflammation in multiple organs by inducing innate immune memory. Specifically, we discovered that IL-1β-mediated epigenetic changes in the myeloid compartment play an unrecognized role in cardiac fibrosis, leading to diastolic dysfunction following ischemic brain injury.

Cardiovascular diseases, such as atrial fibrillation, valvular heart disease, and congestive heart failure, are well-known risk factors for ischemic stroke.^43,44^ However, this relationship is bidirectional, as the incidence of cardiovascular disorders also increases after an initial stroke.^45,46^ After a stroke, more than 60% of patients experience electrocardiographic (ECG) abnormalities,^9^ 25% are diagnosed with serious arrhythmias,^47^ and approximately 19% develop at least one significant cardiac adverse event.^10^ Previous studies in mice have demonstrated that stroke results in chronic systolic dysfunction lasting up to 8 weeks after the brain injury, leading to a delayed reduction in left ventricular ejection fraction and an increase in left ventricular volume.^48,49^ The SICFAIL study, a prospective clinical study involving 696 stroke patients, demonstrated a surprisingly high incidence of cardiac dysfunction after stroke. Diastolic dysfunction was found to be the most prevalent type of cardiac dysfunction, affecting 23% of patients without signs of systolic dysfunction.^50^ This study clearly highlights the previously unrecognized burden of secondary cardiac dysfunction following ischemic stroke.

Although diastolic dysfunction is highly prevalent not only after stroke but also in other comorbidities within the same population (such as smoking, diabetes, and hypertension), it lacks well-defined diagnostic criteria and is not routinely assessed in stroke patients.^51,52^ Recognizing diastolic dysfunction as a pre-symptomatic stage leading to heart failure with reduced ejection fraction could provide an opportunity for targeted secondary prevention if it is identified early.

Patients with chronic systemic inflammatory diseases, such as rheumatoid arthritis, psoriasis, or psoriatic arthritis, are known to have an increased risk of cardiovascular disease. For example, a recent population-based cohort study described an increased incidence of cardiovascular disease in patients with inflammatory bowel disease even in the absence of common risk factors like obesity, lipid disturbances, or hypertension, suggesting that inflammation may be a key factor underlying the development of these cardiovascular complications.^53^ Therefore, it is reasonable to consider that the systemic inflammatory response triggered by the ischemic brain lesion itself may further predispose stroke patients to secondary (inflammatory) vascular events.

Here, we confirmed this hypothesis by identifying that persistent pro-inflammatory changes in myeloid cells after stroke are a causal factor for the development of cardiac fibrosis independent of other predisposing factor for cardiovascular disorders. Innate immune memory—defined as long-term changes in the innate immune cell compartment that alters its responsiveness to a second stimulation—has so far been described in infection models and in vaccination.^13^ Importantly, innate (trained) immunity to a pathogen not only heightened innate immune responsiveness to the same pathogen but was demonstrated to also affect unrelated inflammatory processes, as for example the disease-modifying effect of periodontitis for atherosclerosis progression.^17^ In this study, we make the novel observation of innate immune memory in response to a sterile tissue injury, which links this acute event to the development of a chronic, secondary pathology at a remote organ site.

Our study highlights IL-1β-mediated epigenetic changes and the recruitment of reprogrammed cells to the healthy heart as critical events in the development of chronic secondary organ dysfunction after a stroke. Based on our findings, targeting IL-1β systemically can potentially prevent post-stroke epigenetic changes and the resulting pro-inflammatory effects on remote organ homeostasis. This aligns well with previous research demonstrating IL-1β’s ability to induce innate immune memory and the cardiovascular benefits observed in the CANTOS trial using anti-IL-1β therapy.^37^ However, the trial also revealed significant risks of increased infection rates associated with this non-specific approach that targets a key pro-inflammatory cytokine involved in pathogen clearance. This limitation poses a challenge for the further development of this approach in translational research.

Therefore, in our study, we explored an alternative strategy to limit the recruitment of monocytes from the bone marrow and circulation to remote organs by using a dual chemokine inhibitor targeting CCR2/5 (Cenicriviroc).^38^ Notably, genetic studies have provided evidence of the involvement of CCL2-CCR2 signaling in various cardiovascular disorders, suggesting that targeting this signaling pathway could have beneficial effects on monocyte recruitment.^54^ Moreover, Cenicriviroc has demonstrated safety and efficacy in other conditions, including HIV and hepatosteatosis.^38–40^ Considering our findings regarding the beneficial effects of this approach in limiting secondary organ dysfunction caused by pro-inflammatory reprogrammed circulating monocytes, further clinical evaluation of this therapy in stroke and other sterile tissue injuries seems warranted.

In conclusion, our study provides mechanistic insights into immune-mediated secondary comorbidities following stroke, such as cardiac dysfunction and potentially others. We have identified myeloid innate immune memory as a causal mechanism underlying chronic changes in resident innate immune cells across multiple organs, which contribute to the development or progression of secondary organ dysfunction. Our findings confirm the efficacy of therapeutically targeting this cascade in mice by either preventing epigenetic reprogramming through IL-1β neutralization or by inhibiting the circulation of reprogrammed monocytes to secondary target organs like the heart using CCR2/5 inhibitors. These discoveries offer a novel therapeutic rationale for the secondary prevention of post-stroke comorbidities.

## Acknowledgments

The authors thank Christina Fürle and Kerstin Thuß-Silczak for excellent technical support. The study was supported by the Vascular Dementia Research Foundation, the European Research Council (ERC-StGs 802305, to A.L.) and the German Research Foundation (DFG) under Germany′ s Excellence Strategy (EXC 2145 SyNergy – ID 390857198, to A.L.), and through FOR 2879 (LI-2534/5-1, to A.L).

## Author contributions

Conceptualization, A.S., A.L.; Investigation, A.S., S.Z., J.C., D.M., C.F., O.C.; Formal Analysis, A.S., S.Z., F.C., S.B.G, E.B.; Resources, C.B., K.H., S.F., N.P., Ö.G., E.B., C.S.; Writing – Original Draft, A.S., A.L.; Writing – Review & Editing, A.S., A.L., M.D.; Visualization, A.S., A.L.; Supervision, A.L.; Funding Acquisition, A.L., C.S.

## Declaration of interests

All authors declare no competing interests.

## Notes

### Competing Interest Statement

The authors have declared no competing interest.

