## Supplemental material for "Innate immune memory after brain injury drives inflammatory cardiac dysfunction"

**Supplementary material list:**

- Supplementary Figures S1-S6

Figure S1: Stroke induces long-term inflammatory changes in systemic monocytes/macrophages

Figure S2: Stroke results in chronic cardiac diastolic dysfunction

Figure S3: Bone marrow cellularity and function are chronically altered after stroke

Figure S4: Stroke induces stable innate immune memory

Figure S5: Innate immune memory is mediated by early post-stroke IL-1

Figure S6: Blocking BM-to-heart trafficking of monocytes prevents post-stroke cardiac dysfunction

- STAR Methods
- Supplementary References

**Figure S1**
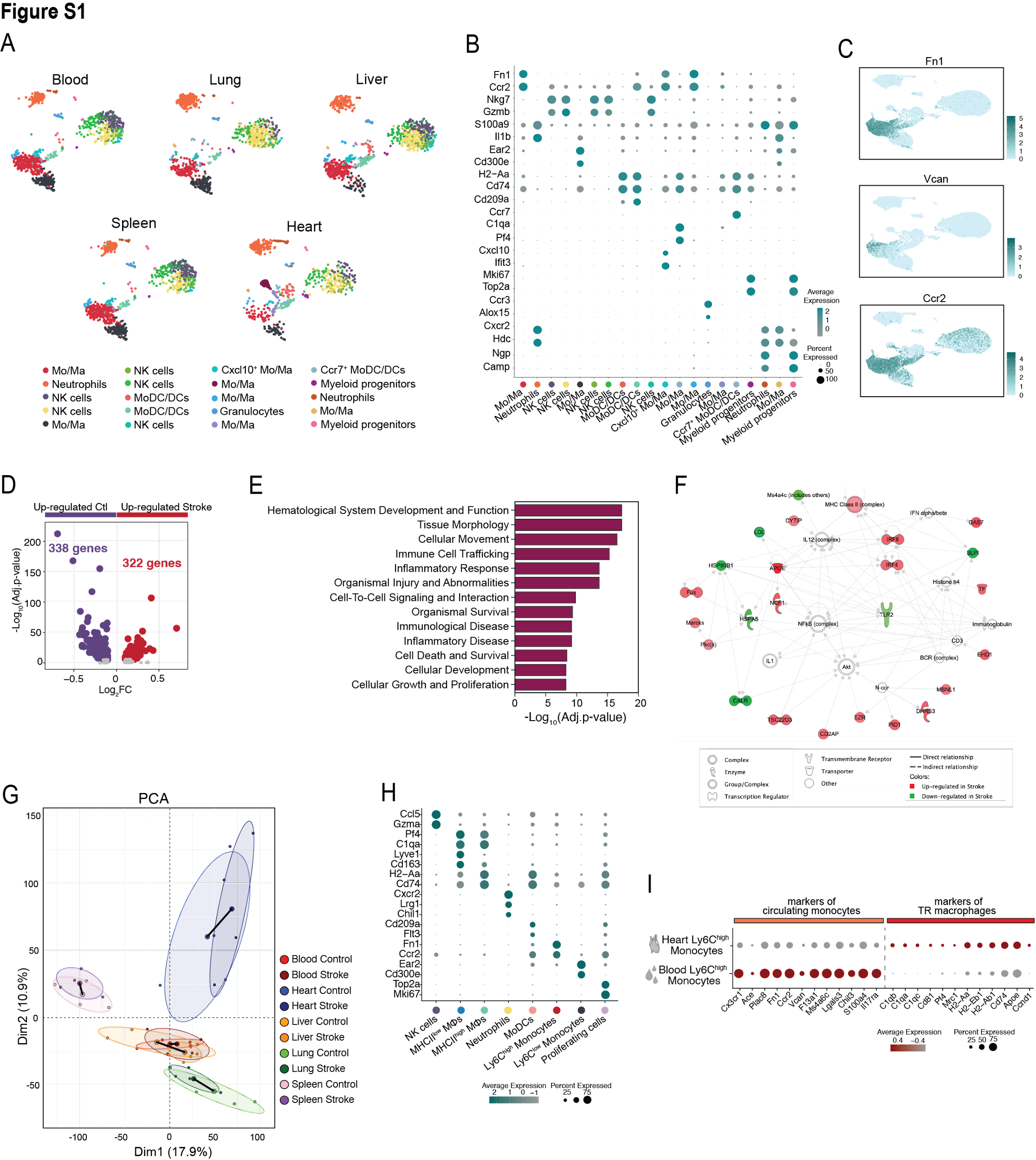

**Figure S1. Stroke induces long-term inflammatory changes in monocytes/macrophages from peripheral organs. (A)** Uniform manifold approximation and projection (UMAP) plots of a representative set of 1,000 CD45^+^ CD11b^+^ cells per peripheral organ, colored by identified populations. **(B)** Dot plot showing the expression profile of core genes for the identification of the populations of CD45^+^ CD11b^+^ cells in peripheral organs. The dot size corresponds to the fraction of cells within each condition expressing the indicated transcript, and the color indicates average expression. **(C)** UMAP plots with expression overlay of *Fn1*, *Vcan* and *Ccr2* genes, highly expressed in the selected populations of monocyte/macrophages. **(D)** Volcano plot showing the up (322 genes, red) and down-regulated genes (338 genes, purple) between the 2 selected populations of monocyte/macrophages from stroke and control mice. Colored genes are p<0.05 and |fold-change|>1.25. **(E)** Pathway analysis was performed using Ingenuity Pathway Analysis (IPA, Qiagen) using the DEG from the 2 selected populations of monocytes/macrophages, with an adjusted p value <0.05 and |fold-change|>1.58. Top diseases and functions categories sorted by p value are displayed. **(F)** Functional gene interaction network analysis using IPA. Genes are colored based on fold-change values determined by the single cell mRNA sequencing analysis, where red indicates an increase in stroke and green in control animals. **(G)** PCA plot displaying all analyzed samples from all peripheral organs. The PCA was calculated from a total of 18,834 genes identified in peripheral CD45^+^CD11b^+^ myeloid cells. The Euclidian distances between stroke and control clusters per organ are indicated in black lines. **(H)** Dot plot showing the expression profile of selected genes key for the identification of the cell populations in the heart. The dot size corresponds to the fraction of cells within each condition expressing the indicated transcript, and the color indicates average expression. **(I)** Dot plot showing the expression levels of selected gene features in Ly6C^high^ monocytes from the heart and blood. Genes were selected from the list of DEG between both cell populations. The dot size corresponds to the fraction of cells within each condition expressing the indicated transcript, and the color indicates average expression.

**Figure S2**

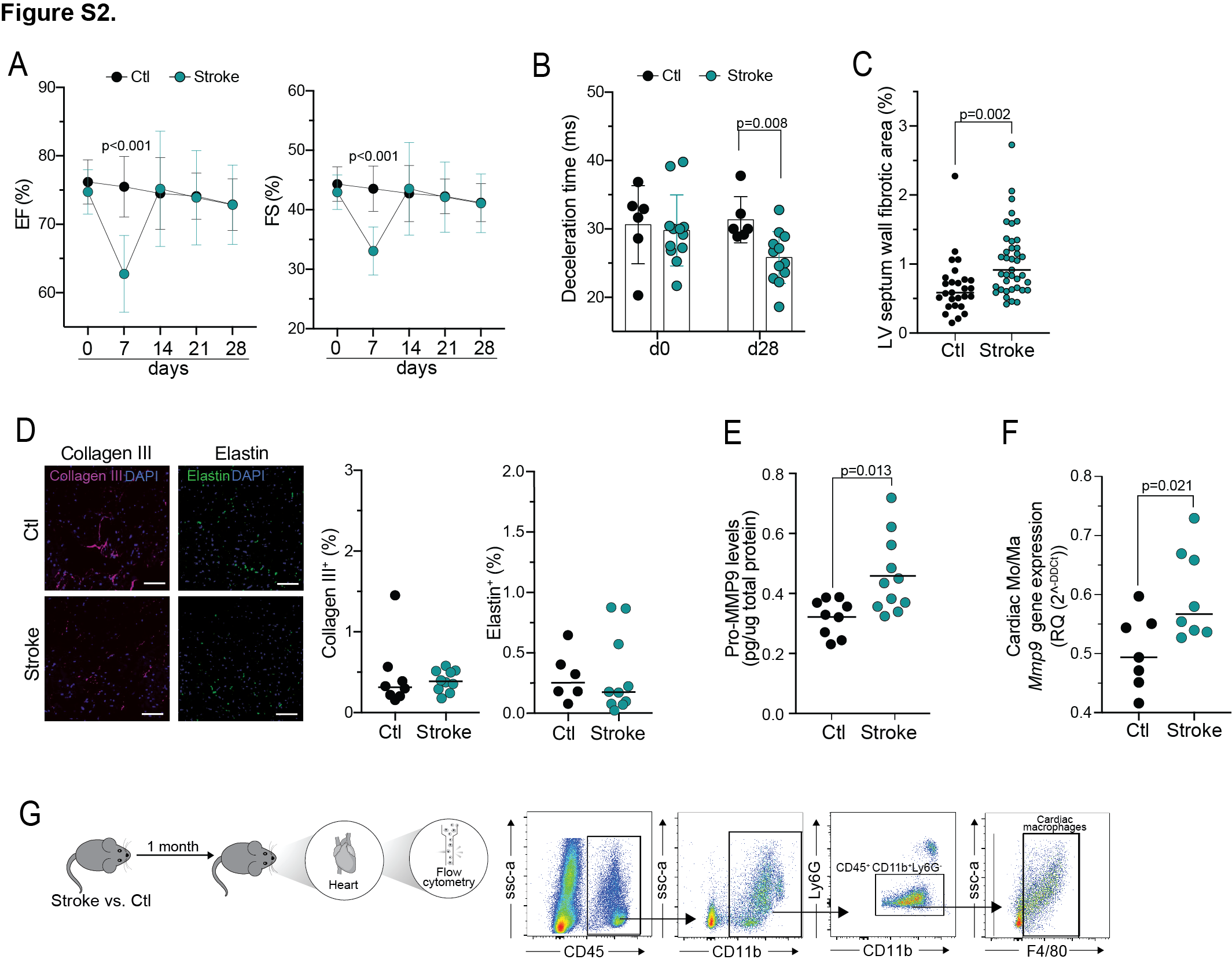

**Figure S2. Stroke results in chronic cardiac diastolic dysfunction. (A)** Quantification of the ejection fraction (EF, left panel) and fractional shortening (FS, right panel) at indicated time points (days) before (day 0) and after stroke, and control (U tests, n=6/12 per group). **(B)** Quantification of the deacceleration time at day 0 and 28 in control and stroke mice (U tests, n=6/12 per group). **(C)** Quantification of cardiac fibrosis in the LV septum wall (t test, n=7/10 per group, 4 heart sections per mouse). **(D)** Representative images of immunofluorescence staining for the detection of collagen III and elastin in heart coronal sections (left; scale bar = 50 µm). Corresponding quantification of collagen III and elastin content, expressed in percentage of total area of the LV free wall (right; t test, n = 8/10 per group). **(E)** Total pro-MMP9 protein levels were measured in heart samples from stroke and control mice. Pro-MMP9 protein levels are normalized to total protein content (U-test, n=9/11 per group). **(F)** RT-qPCR was performed on sorted monocyte/macrophages (Mo/Ma, CD45^+^Ly6G^-^CD11b^+^**)** to measure the expression levels of *Mmp9* mRNA, quantified relative to the expression of the housekeeping gene encoding for *Ppia* and normalized to control levels (U test, n=7 per group). **(G)** Schematic experimental design: cardiac myeloid cells were isolated from the heart of stroke and control mice one month after stroke and analyzed by flow cytometry. Representative gating strategy for the non-neutrophil cardiac myeloid cells (CD45^+^ Ly6G^-^ CD11b^+^) and cardiac macrophages (CD45^+^ Ly6G^-^ CD11b^+^F4/80^+^).

**Figure S3**

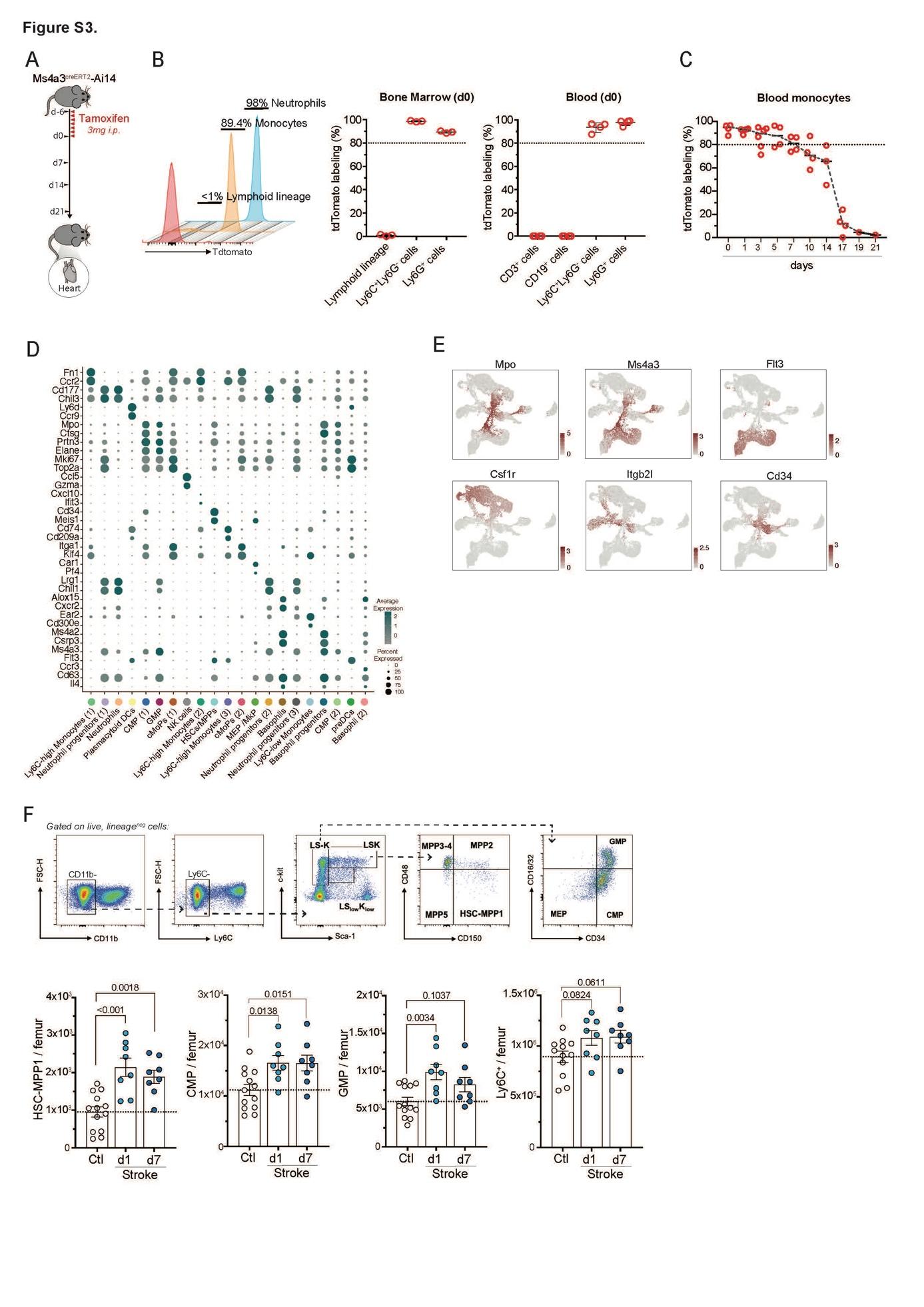

**Figure S3. Bone marrow cellularity and function are chronically altered after stroke. (A)** Schematic experimental design: Ms4a3^creERT2^xAi14 mouse were treated with tamoxifen for 7 consecutive days. Mice were sacrificed at different time points within the 3 weeks after tamoxifen treatment and the heart myeloid cells were analyzed by flow cytometry (left panel). **(B)** TdTomato labeling of lymphocytes (CD3+, CD4+, CD8a+, CD19+, Ter119+), monocytes (Ly6C+Ly6G-) and neutrophils (Ly6C+Ly6G+) in bone marrow (mid panels) and blood (right panel) after 7 daily doses of tamoxifen (n=3/4). **(C)** TdTomato labeling of monocytes (Ly6C+Ly6G-) in blood over time after 7 daily doses of tamoxifen (n=4). **(D)** Dot plot showing the expression profile of selected genes key for the identification of the populations of lymphoid (CD3, CD4, CD8a, CD19 and Ter119)-lineage and neutrophil (Ly6G) negative myeloid cells sorted from the BM of control and stroke mice 1 month after stroke (n=4/group). The dot size corresponds to the fraction of cells within each condition expressing the indicated transcript, and the color indicates average expression. **(E)** UMAP plots with expression overlay of *Mpo*, *Ms4a3, Csf1rm Itgb2l, Cd34* and *Flt3* genes. **(F)** Representative gating strategy for BM progenitor cells (upper panel). Corresponding cell count quantifications of hematopoietic stem cells-multipotent progenitors 1 (HSC-MPP1), common myeloid progenitors (CMP), granulocyte-monocyte progenitors (GMP) and Ly6C^+^ monocytes in stroke and control mice at 1 and 7 days after stroke (bottom panels) (U-test; n=9/13 per group). LSK: Lin-Sca1+c-Kit+; LS-K: Lin-Sca1-c-Kit+; LS^low^K^low^: Lin-Sca1^low^c-Kit^low^; MMP: multipotent progenitors.

**Figure S4**

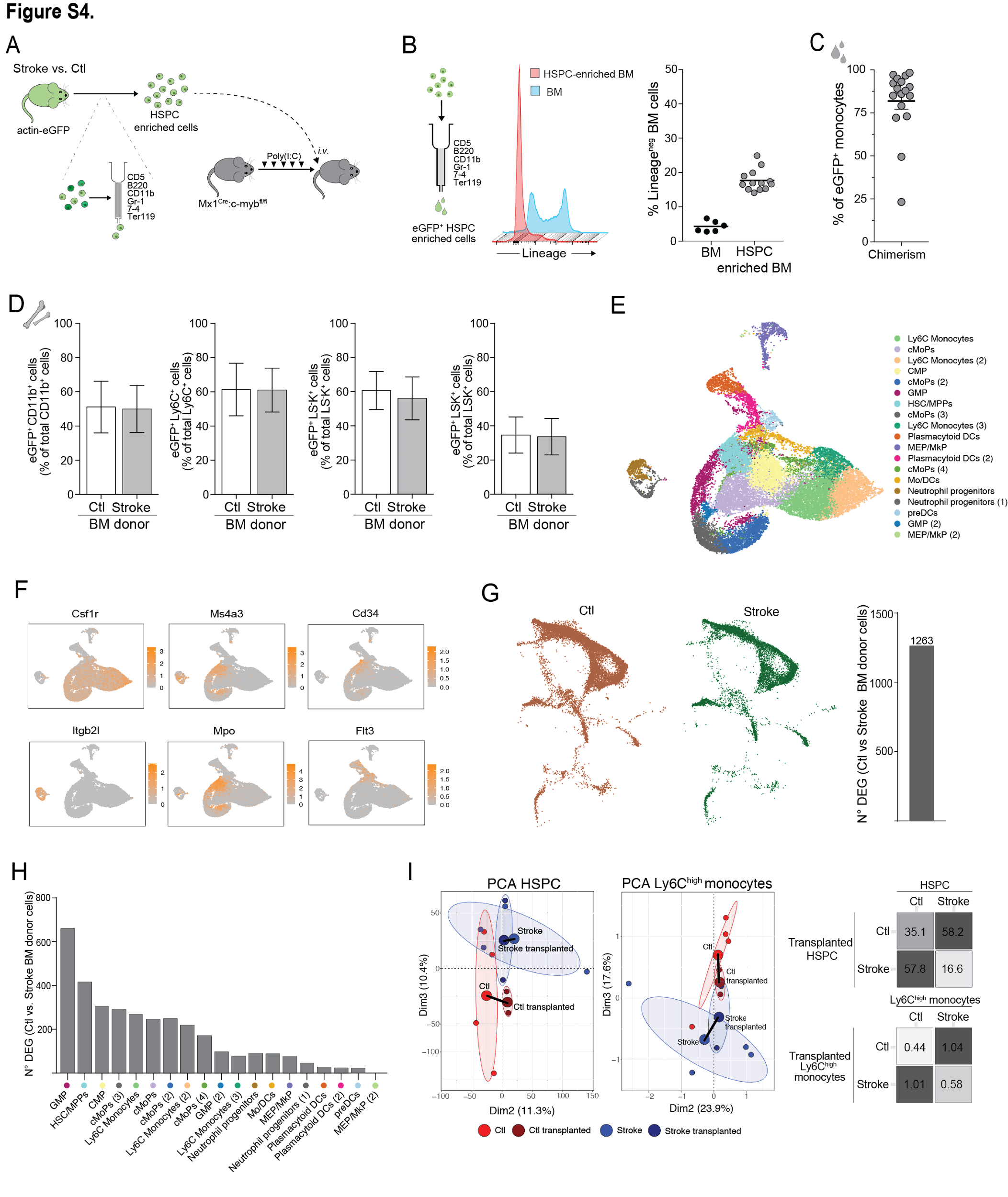

**Figure S4. Stroke induces stable innate immune memory. (A)** Schematic experimental design: bone marrow (BM) cells were isolated from actin-eGFP control and stroke mice 1 month after stroke and enriched for hematopoietic stem and progenitor cells (HSPC). HSPC-enriched cells were transplanted into BM-depleted Mx1^Cre^:c-myb^fl/fl^ mice. **(B)** Schematic of the enrichment of BM cells for HSPC by negative selection of lineage markers (CD5, B220, CD11b, Gr-1, 7-4, Ter119, left panel). Histogram of cells before and after HSPC enrichment showing the proportion of lineage^neg^ and lineage^pos^ BM cells (mid panel). Corresponding flow cytometry quantification of the percentage of lineage^neg^ cells before and after HSPC enrichment (n=6/12 per group, right panel). **(C)** Blood chimerism 1 month after BM transplantation, depicted as the percentage of eGFP^+^ monocytes in blood from recipient mice. **(D)** Percentage of eGFP^+^ donor-derived CD11b^+^, Ly6C^+^, LS-K^+^ (Lineage^-^ Sca^-^ c-kit^+^) and LSK^+^ (Lineage^-^ Sca^+^ c-kit^+^) cells from recipient mice (n=6 per group). Results indicate no difference in repopulation efficacy between control and stroke donors. **(E)** UMAP plot of 25,358 myeloid cells from the BM of transplanted mice, colored by identified populations. **(F)** UMAP plots with expression overlay of *Mpo*, *Ms4a3, Csf1rm Itgb2l, Cd34* and *Flt3* genes. **(G)** UMAP plot of myeloid cells from the BM of transplanted mice, split by condition (left panel) and number of differentially expressed genes (DEG) between stroke and control (adjusted p value <0.05). **(H)** Number of DEG between stroke and control conditions per cell type. **(I)** PCA plot displaying individual samples from the BM of stroke (light blue) and control mice (light red) and the BM of recipient mice transplanted with stroke (dark blue) and control (dark red) eGFP^+^ HSPC-enriched BM cells (left panels). The PCA was calculated from a total of 18,834 genes identified in BM myeloid cells. The Euclidian distances between stroke samples and control samples are indicated in black lines and were calculated from the mean of each cluster.

**Figure S5**

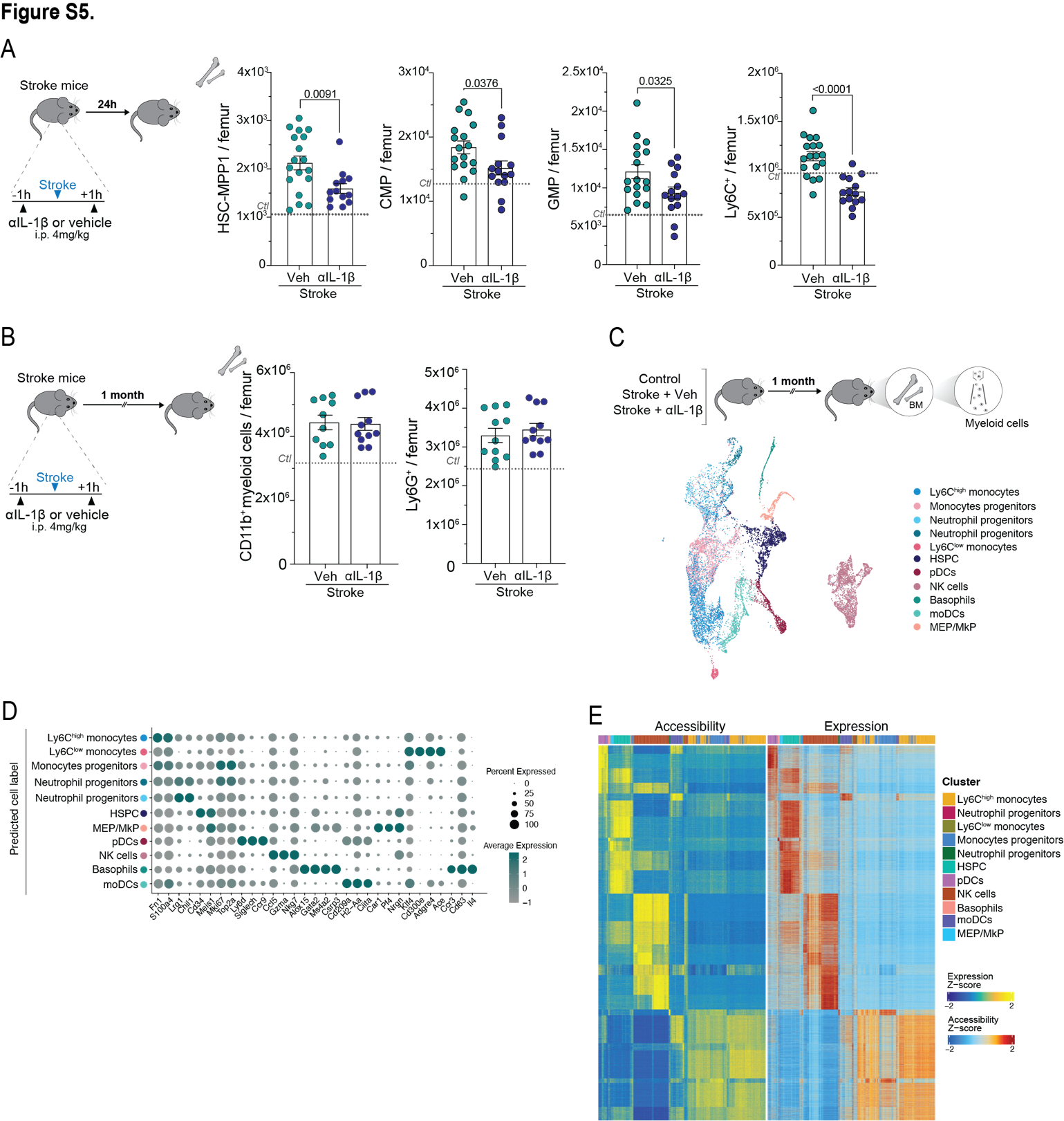

**Figure S5. Innate immune memory is mediated by early post-stroke IL-1β. (A)** Schematic experimental design: mice received Interleukin (IL)-1β neutralizing antibodies or vehicle 1h before and 1h after stroke induction. Twenty-four hours later, bone marrows (BM) were collected for flow cytometry (left panel). Cell count quantifications of hematopoietic stem cells-multipotent progenitors 1 (HSC-MPP1), common myeloid progenitors (CMP), granulocyte-monocyte progenitors (GMP) and total Ly6C^+^ monocytes in BM from anti-IL1β or vehicle-treated stroke mice (U-test; n=13/18 per group). **(B)** Same experimental design as in (A), but BM were collected 1 month after stroke for flow cytometric cell count quantifications of total CD11b^+^ myeloid cells and Ly6G^+^ neutrophils (U-test; n=10/11 per group). **(C)** Schematic experimental design: nuclei were isolated from lineage-negative myeloid cells sorted from the BM of control and anti-IL1β or vehicle-treated stroke mice 1 month after stroke (n=3/group). The transcriptomic profiles of sorted nuclei were analyzed using single-nuclei ATAC-sequencing (upper panel). UMAP plot of the 13,520 myeloid nuclei sorted from the BM of control and anti- IL1β and vehicle-treated stroke mice, colored by the transferred cell labels from the single cell mRNA sequencing dataset. **(D)** Dot plot showing the expression profile of selected genes key for the identification of the populations based on the single cell mRNA sequencing data. The dot size corresponds to the fraction of nuclei within each condition expressing the indicated transcript, and the color indicates average expression. **(E)** Side-by-side heatmaps showing the correspondence links of peaks (left) and gene (right) from the single nuclei ATAC sequencing and single cell mRNA sequencing, respectively.

**Figure S6**

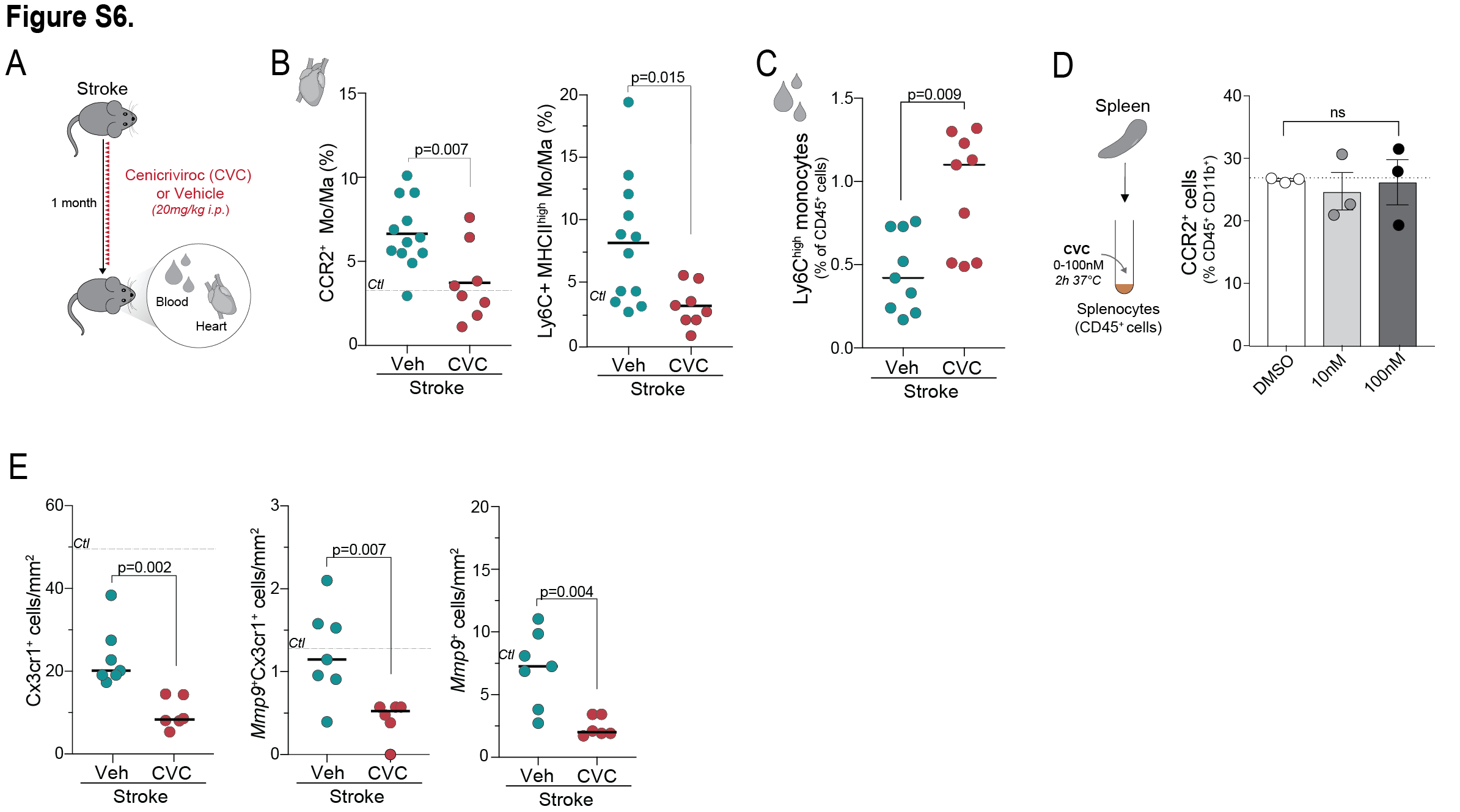

**Figure S6. Blocking BM-to-heart trafficking of monocytes prevents post-stroke cardiac dysfunction. (A)** Schematic experimental design: Stroke mice were administered daily with the dual C-C chemokine receptors type 2 and 5 antagonist Cenicriviroc (Stroke+CVC) or vehicle (Stroke) for 28 days. Mice were sacrificed at day 28 and the heart and blood were collected for flow cytometry and histological analysis. **(B)** Flow cytometry of total CCR2^+^ and Ly6C^+^MHC-II^high^ monocytes/macrophages in the heart of stroke mice treated with CVC or vehicle (t test; n=8/12 per group). **(C)** Flow cytometry for circulating Ly6C^high^ monocytes in CVC- or vehicle-treated stroke mice, expressed as percentage of total circulating CD45^+^ cells. Dashed lines indicate mean values in control mice (t test; n=9 per group). **(D)** Schematic experimental design: splenocytes were freshly isolated from naïve mice and treated with CVC at a dose of 0 (DMSO), 10 and 100 nM. Two hours after treatment, the percentage of live CCR2+ cells was evaluated by flow cytometry. **(E)** smFISH quantification for the detection of total *Cx3cr1*^+^ cells (left panel), *Mmp9*^+^ *Cx3cr1*^+^ cells (mid panel) and *Mmp9*^+^ cells (right panel) in the hearts of CVC- or vehicle-treated stroke mice. DAPI was used as nuclear dye. Dashed lines indicate mean values in control mice (t test, n=6/7 per group).

**STAR METHODS**

**KEY RESOURCES TABLE**

| REAGENT or RESOURCE | SOURCE | IDENTIFIER |
| --- | --- | --- |
| Antibodies |  |  |
| Anti-mouse CD45 (30-F11), Fluor™ 450 | eBioscience | Cat# 48-0451-82 |
| Anti-mouse CD16/CD32 (2.4G2), BV480 | BD Biosciences | Cat# 746324 |
| Anti-mouse CD11b (M1/70), FITC | BioLegend | Cat# 557396 |
| Anti-mouse Ly6C (HK1.4), Brilliant Violet 570™ | BioLegend | Cat# 128029 |
| Anti-mouse Ly6G (1A8-Ly6g), PE-eFluor™ 610 | ThermoFisher Scientific | Cat# 61-9668-82 |
| Anti-mouse Sca1 (D7), Brilliant Violet 421™ | BioLegend | Cat# 108127 |
| Anti-mouse CD115 (AFS98), BV711 | BD Biosciences | Cat# 750890 |
| Anti-mouse CD150 (TC15-12F12.2), PE | BioLegend | Cat# 115903 |
| Anti-mouse *ckit* (CD117) (2B8), PE-Cy™7 | BD Biosciences | Cat# 558163 |
| Anti-mouse CD135 (A2F10), APC | BioLegend | Cat# 135309 |
| Anti-mouse CD11c (N418), Brilliant Violet 650™ | BioLegend | Cat# 117339 |
| Anti-mouse CD127 (A7R34), Brilliant Violet 785™ | BioLegend | Cat# 135037 |
| Anti-mouse CD48 (HM48-1), PerCP | Elabscience | Cat# E-AB-F1017UF |
| Anti-mouse CD34 (RAM34), Alexa Fluor® 700 | BD Biosciences | Cat# 560518 |
| Anti-mouse CD3 (17A2), APC-eFluor™ 780 | eBioscience | Cat# 47-0032-80 |
| Anti-mouse CD4 (RM4-5), APC-eFluor™ 780 | Invitrogen | Cat# 47-0042-82 |
| Anti-mouse CD8a (53-6.7), APC/Cyanine7 | BioLegend | Cat# 100714 |
| Anti-mouse CD19 (eBio1D3 (1D3)), APC-eFluor™ 780 | Invitrogen | Cat# 47-0193-82 |
| Anti-mouse TER-119/Erythroid Cells (TER-119), APC-Cy™7 | BD Biosciences | Cat# 560509 |
| Anti-mouse CD16/CD32() | Invitrogen | Cat# 14-0161-86 |
| Anti-mouse CD11b (93), APC-Cyanine7 | eBioscience | Cat# A15390 |
| Anti-mouse Ly6C (HK1.4), PerCP/Cyanine5.5 | BioLegend | Cat# 128011 |
| Anti-mouse F4/80 (BM8), PE-Cyanine7 | eBioscience | Cat # 25-4801-82 |
| Anti-mouse CCR2 (SA203G11) (Brilliant Violet 785™) | BioLegend | Cat # 150621 |
| Anti-mouse MHC Class II (NIMR-4), PE | eBioscience | Cat # 12-5322-81 |
| Anti-mouse CCR2 (SA203G11), APC | BioLegend | Cat # 150628 |
| Anti-mouse CD45 (30-F11), APC-Cy7 | BioLegend | Cat # 103116 |
| Anti-mouse CD11b (M1/70), BV510 | BD Biosciences | Cat #562950 |
| Anti-mouse Collagen I | Invitrogen | Cat # PA5-95137 |
| Anti-mouse Collagen III | Abcam | Cat # ab184993 |
| Anti-mouse Elastin | ThermoFisher Scientific | Cat # BS-1756R |
| Anti-GFP Nanobody conjugated to Alexa Fluor® 488 (GFP-booster) | ChromoTek | Cat# gb2AF488 |
| Anti-mouse Hashtag 1 Antibody TotalSeq-B0301 (M1/42) | BioLegend | Cat# 155831 |
| Anti-mouse Hashtag 2 Antibody TotalSeq-B0302 (M1/42) | BioLegend | Cat# 155833 |
| Anti-mouse Hashtag 3 Antibody TotalSeq-B0303 (M1/42) | BioLegend | Cat# 155835 |
| Anti-mouse Hashtag 4 Antibody TotalSeq-B0304 (M1/42) | BioLegend | Cat# 155837 |
| ***Continued*** |  |  |
| REAGENT or RESOURCE | SOURCE | IDENTIFIER |
| Anti-mouse Hashtag 5 Antibody TotalSeq-A0305 (M1/42) | BioLegend | Cat# 155809 |
| Anti-human CD14 | NovusBio | Cat# NBP2-67630 |
| Anti-human CCR2 | R&D systems | Cat# MAB150 |
| Goat Anti-Rabbit (H+L) Crossed-adsorbed Secondary antibody | Abcam | Cat# 214880 |
| Alexa Flour 647 goat anti-rabbit secondary antibody | Invitrogen | Cat # A-21247 |
| Anti-mouse/rat IL-1β (InVivoMAb) | BioXcell | Cat # BE0246 |
| Critical commercial assays |  |  |
| Picro-Sirius Red Stain Kit (Cardiac Muscle) | Abcam | Cat# ab245887 |
| Pierce™ BCA Protein Assay Kit | ThermoFisher | Cat# 23227 |
| Colloidal Blue Staining Kit | Invitrogen | Cat# LC6025 |
| High-Capacity cDNA Reverse Transcription Kit | Applied Biosystems | Cat# 4368814 |
| QuantiTect SYBR® Green PCR Kits | Qiagen | Cat# 204143 |
| RNAscope® Multiplex Fluorescent Detection Kit v2 | ACDbio | Cat# 323110 |
| RNAscope® TSA Buffer Pack | ACDbio | Cat# 322810 |
| TSA Plus Cyanine 3 (Cy3) detection Kit | PerkinElmer | Cat# NEL744001KT |
| TSA Plus Cyanine 5 (Cy5) detection kit | PerkinElmer | Cat# NEL745001KT |
| pro-MMP-9 Mouse ELISA Kit | ThermoFisher Scientific | Cat# EMMMP9 |
| Chromium Next GEM Chip G Single Cell Kit | 10x Genomics | Cat# 1000127 |
| Chromium Next GEM Chip H Single Cell Kit | 10x Genomics | Cat# 1000162 |
| Chromium Next GEM Single Cell ATAC Kit v2 | 10x Genomics | Cat# 1000406 |
| 3ʹ Feature Barcode Kit | 10x Genomics | Cat# 1000262 |
| Library Construction Kit | 10x Genomics | Cat# 1000190 |
| Single Index Kit T Set A | 10x Genomics | Cat# 1000213 |
| Single Index Kit N Set A | 10x Genomics | Cat# 1000212 |
| Lineage Cell Depletion Kit, mouse | Miltenyi Biotec | Cat# 130-090-858 |
| MACSprep^TM^ Chimerism CD34 MicroBead Kit human | Miltenyi Biotec | Cat# 130-120-673 |
| Click-iT™ EdU Alexa Fluor™ 647 Flow Cytometry Assay Kit | Invitrogen | Cat# C10419 |
| Agilent High Sensitivity DNA kit | Agilent | Cat# 5067-4626 |
| Stainless steel beads 5mm | Qiagen | Cat# 69989 |
| Biological samples |  |  |
| Human heart paraffin sections | NCBN (Japan) | (Please see details in Methods section) |
| Human heart paraffin blocks | LMU (Munich) | (Please see details in Methods section) |
| Chemicals, peptides, and recombinant proteins |  |  |
| Cenicriviroc | Biorbyt | Cat# orb402001 |
| Kolliphor | Sigma Aldrich | Cat# 5135 |
| Murine recombinant IL-1beta | R&D systems | Cat# 401-ML |
| 5-ethynyl-2´-deoxyuridine (EdU) | Invitrogen | Cat# E10187 |
| Tamoxifen | Sigma Aldrich | Cat# 85256 |
| Poly(I:C) HMW | InvivoGen | Cat# tlrl-pic |
| Miglyol 812 | Caelo | Cat# 3274 |
| ***Continued*** |  |  |
| REAGENT or RESOURCE | SOURCE | IDENTIFIER |
| Collagenase type XI | Sigma Aldrich | Cat# C7657 |
| Collagenase type I | Sigma Aldrich | Cat# SCR103 |
| Deoxyribonuclease I | Sigma Aldrich | Cat# 9003-98-9 |
| Hyaluronidase | Sigma Aldrich | Cat# 37326-33-3 |
| Histopaque®-1077 | Sigma Aldrich | Cat# 10771 |
| Zombie NIR | BioLegend | Cat# 423105 |
| Propidium Iodide | eBiosciences | Cat# 00-6990-42 |
| 7-AAD (7-Aminoactinomycin D) | ThermoFisher Scientific | Cat# A1310 |
| BD Horizon™ Brilliant Stain Buffer | BD Biosciences | Cat# 563794 |
| Cell Staining Buffer | BioLegend | Cat# 420201 |
| Fast Green FCF (0.1 %) | Morphisto | Cat# 16596 |
| Eosin Y solution (1%) | Carl Roth | Cat# 17372-87-1 |
| Mayer's hematoxylin solution | Sigma Aldrich | Cat# MHS32 |
| Acetic acid | Carl Roth | Cat# 64-19-7 |
| Ethanol | SAV Liquid | Cat# 64-17-5 |
| Paraformaldehyde | Morphisto | Cat# 11762.00100 |
| Agarose | VWR Chemicals | Cat# 9012-36-6 |
| Goat serum | ThermoFisher Scientific | Cat# 16210064 |
| Bovine Serum Albumin | Sigma Aldrich | Cat# 9048-46-8 |
| Gelatin from cold water fish skin | Sigma Aldrich | Cat# 9000-70-8 |
| DAPI (4’-6-Diamidino-2-Phenylindole-dihydrochloride) | ThermoFisher Scientific | Cat# D3571 |
| Eukitt® Quick-hardening mounting medium | Sigma Aldrich | Cat# 25608-33-7 |
| Protease and Phosphatase Inhibitors | ThermoFisher Scientific | Cat# A32959 |
| RIPA lysis/extraction buffer | ThermoFisher Scientific | Cat# 89900 |
| Zymogram Plus (Gelatin) gels 10% | Invitrogen | Cat# ZY00102BOX |
| Zymogram Renaturing Buffer (10X) | Invitrogen | Cat# LC2670 |
| Zymogram Developing Buffer (10X) | Invitrogen | Cat# LC2671 |
| Novex™ Tris-Glycin SD-Probenpuffer (2X) | Invitrogen | Cat# LC2676 |
| RNAscope® Probe- Mm-Mmp9-C2 | ACDbio | Cat# 315941-C2 |
| RNAscope® Probe- Mm-Cx3cr1 | ACDbio | Cat# 314221 |
| RNAscope® Probe- Hs-MMP9 | ACDbio | Cat# 311331 |
| RNAscope® Probe- Hs-CD14 | ACDbio | Cat# 418808-C2 |
| RNAscope® Wash Buffer Reagents | ACDbio | Cat# 310091 |
| RNAscope® Target Retrieval Reagents | ACDbio | Cat# 322000 |
| Tween-20 | BioRad | Cat# 1662404 |
| Digitonin 5% | ThermoFisher Scientific | Cat# BN2006 |
| Nuclei EZ Lysis Buffer | Sigma Aldrich | Cat# NUC-01 |
| Dulbecco's Phosphate Buffered Saline | Sigma Aldrich | Cat# D8537-500ML |
| Fetal calf serum (FBS) | GIBCO | Cat# 105000-064 |
| Buffer PKD | Qiagen | Cat# 1034963 |
| Proteinase K | Qiagen | Cat# 19131 |
| Dynabeads™ Oligo(dT)25 | ThermoFisher Scientific | Cat# 61005 |
| Recombinant RNase Inhibitor | Takara | Cat# 2313A |
| UltraPure™ SSPE, 20X | ThermoFisher Scientific | Cat#15591043 |
| Experimental models: Organisms/strains |  |  |
| Mouse: C57BL6/J | Charles River | Strain #: 000664 |
| Mouse: eGFP (C57BL/6-Tg(CAG-EGFP)131Osb/LeySopJ) | JAX | Strain #: 006567 |
| Mouse: Mx1Cre (B6.Cg-Tg(Mx1-cre)1Cgn/J) | JAX | Strain #: 003556 |
| Mouse: Myb^fl/fl^ (B6.129P2-Mybtm1Cgn/TbndJ) | JAX | Strain #: 028881 |
| Mouse: Ai14 (B6.Cg-Gt(ROSA)26Sortm14(CAG-tdTomato)Hze/J) | JAX | Strain #: 007914 |
| Mouse: Ms4a3(creERT2) | This paper | N/A |
| Oligonucleotides |  |  |
| Mmp9  Forward 5' GCT CCT GGC TCT CCT GGC TT 3' Reverse 5' GTC CCA CCT GAG GCC TTT GA 3' | Metabion | N/A |
| Ppia  Forward 5' ACA CGC CAT AAT GGC ACT GG 3' Reverse 5' ATT TGC CAT GGA CAA GAT GC 3' | Metabion | N/A |
| Software and algorithms |  |  |
| FlowJo v.10.6 | Treestar Inc. | N/A |
| GraphPad Prism 9 | Graphpad Inc. | N/A |
| Image J 1.53c | NIH | N/A |
| ZEN | ZEISS | N/A |
| LAS X Office | Leica | N/A |
| Vevo LAB 5.5.0 | FUJIFILM Visual Sonics | N/A |
| Cell Ranger v.7.1.0 | 10x Genomics | N/A |
| Cell Ranger ATAC v.2.1.0 | 10x Genomics | N/A |
| R package Seurat 4.2.0 | ^1^ | N/A |
| R package Signac 1.9.0 | ^2^ | N/A |
| Monocle3 v.1.3.1 | ^3^ | N/A |
| R package ArchR v.1.0.1 | ^4^ | N/A |
| FastQC | ^5^ | N/A |
| STAR | ^6^ | N/A |
| Cutadapt | ^7^ | N/A |
| featureCounts | ^8^ | N/A |
| DESeq2 | ^9^ | N/A |
| Ingenuity Pathway Analysis | QIAGEN | N/A |
| g:Profiler | ^10^ | N/A |
| Deposited data |  |  |
| Raw single cell mRNAseq data | This paper | GSE232098 |
| Raw single nuclei ATACseq data | This paper | GSE230692 |
| Raw bulk mRNAseq data | This paper | GSE232550 |
| Others |  |  |
| Silicone rubber-coated monofilament | Doccol | Cat# 602223PK10Re |
| ALZET micro-osmotic pumps | Alzet | Cat#1007D |

**RESOURCE AVAILABILITY**

**Lead contact**

**Materials availability**

This study did not generate new unique reagents.

**Data and code availability**

Single-cell RNA-seq and single-nuclei ATAC-seq data have been deposited at GEO and are publicly available as of the date of publication. Accession numbers are also listed in the key resources table. This paper does not report original code. The used scripts for bioinformatic analyses of sequencing data are available at <https://github.com/Lieszlab>.

**EXPERIMENTAL MODEL AND SUBJECT DETAILS**

**Clinical patient population**

Human postmortem material was obtained from the National Center Biobank Network (NCBN) from Japan and the Institute of Legal Medicine, LMU University from Munich. Ethical approval for the use of human postmortem material was granted according to institutional ethics board protocol and national regulations.

|  | Stroke (n=12) | Control (n=7) |
| --- | --- | --- |
| Age, *years* | 79 (13.5) | 80 (26.0) |
| Sex (male), *n (%)* | 75% (9) | 42.9% (3) |
| Baseline NIHSS | 21 (17.3) | N/A |
| Time from stroke to death, *days* | 50 (42.0) | N/A |

*Shown as Median (Interquartile range, IQR), unless indicated*

**Animal experiments**

All animal procedures were performed in accordance with the guidelines for the use of experimental animals and were approved by the respective governmental committees (Regierungspraesidium Oberbayern, the Rhineland Palatinate Landesuntersuchungsamt Koblenz). Wild-type C57BL6/J mice were purchased from Charles River. eGFP reporter mice (C57BL/6-Tg (CAG-EGFP)131Osb/LeySopJ) and Ai14 reporter mice (B6.Cg-Gt(ROSA)26Sortm14(CAG-tdTomato)Hze/J) were purchased from the Jackson Laboratory (US), and bred and housed at the animal core facility of the Center for Stroke and Dementia Research (Munich, Germany). The Mx1-Cre mice (B6.Cg-Tg(Mx1-cre)1Cgn/J) and the Myb^fl/fl^ floxed mutant mice (B6.129P2-Mybtm1Cgn/TbndJ) were also purchased from Jackson Laboratory (US), and Mx1^Cre^:*c-Myb*^fl/fl^ mice were generated, bred and housed at the animal core facility of the Walter-Brendel-Center for Experimental Medicine (Munich, Germany). Ms4a3^creERT2^ mice (B6J-Ms4a3em1(CreERT2)-Gt(ROSA)26Sortm14(CAG-tdTomato)) were generated at the animal core facility of the Center for Stroke and Dementia Research (Munich, Germany), by CRISPR-Cas9-mediated insertion of an IRES-CreERT2 cassette into the 3’ un-translated region (3’UTR) of the *Ms4a3* gene in C57BL/6 zygotes. Genotyping was done by PCR using the following primers: Ms4a3 forward primer 5’- GACATTGCAGACGGGATGTAT-3’; Ms4a3 reverse primer 5’- ATCCATGGAGGTGTCATAGACCA-3’; CreERT2 forward primer 5’-AACACCCCGTGAAACTGCTC-3’.

All mice had free access to food and water at a 12 h dark-light cycle. Data were excluded from all mice that died during surgery. Animals were randomly assigned to treatment groups and all analyses were performed by investigators blinded to group allocation. All animal experiments were performed and reported in accordance with the ARRIVE guidelines.^11^

**METHOD DETAILS**

**Transient proximal cerebral artery occlusion model**

Transient intraluminal occlusion of the middle cerebral artery (MCA) was performed as previously described.^12^ Briefly, mice were anaesthetized with isoflurane delivered in 100% O_2_. A midline neck incision was made and the common carotid artery and left external carotid artery were isolated and ligated; a 2-mm silicon-coated filament was introduced via a small incision in the external carotid artery and advanced towards the internal carotid artery, therefore occluding the MCA. After 60 min of occlusion, the animals were re-anesthetized, and the filament was removed. MCA occlusion and reperfusion were confirmed by the corresponding decrease or increase in the blood flow, respectively, measured by a laser Doppler probe affixed to the skull above the MCA territory (decrease in the laser Doppler flow signal >80% of baseline value and increase in the laser Doppler flow signal>80% of baseline value before reperfusion). After recovery, the mice were kept in their home cage with facilitated access to water and food. Body temperature was maintained at 37°C throughout surgery using a feedback-controlled heating pad. The overall mortality rate of stroke mice was approximately 30 %. Exclusion criteria: 1. Insufficient MCA occlusion (a reduction in blood flow to >20% of the baseline value); 2. Insufficient MCA reperfusion (an increase in blood flow of >80% of the baseline value before removing the filament); 3. Death during the surgery.

**Drug administrations**

**5-ethynyl-2´-deoxyuridine (EdU)**

Mice received one i.p. injection of EdU dissolved in 0.9% Sodium Chloride 4 h prior to euthanasia, at a dose of 5 mg kg^-1^ body weight, in a final volume of 250 μl. For continuous and controlled EdU delivery, osmotic pumps (ALZET) containing 100 μl of EdU at a concentration of 10 mg/ml were implanted i.p. into mice for 1 week.

**Anti-IL-1β**

Mice received two i.p. injections of antagonizing anti-IL-1β or vehicle (0.9% NaCl) 1 h prior to and 1 h after surgery, at a dose of 4 mg kg^-1^ body weight, in a final volume of 200 μl.

**CCR2/CCR5 antagonist Cenicriviroc (CVC)**

Mice received daily i.p. injections of CVC or vehicle (30% Kolliphor and 70% 0.9% NaCl) at a dose of 20 mg kg^-1^ body weight, in a final volume of 250 μl.^13^ First dose was given after stroke induction and then daily for 28 days.

**Tamoxifen**

Tamoxifen was prepared by dissolving in Miglyol 812 for a final concentration of 20 mg/mL and stored at 4 ºC. Mice received daily i.p. injections of Tamoxifen solution at a dose of 120 mg kg^-1^ body weight, in a final volume of 150 μl, for 7 consecutive days.

**Echocardiography**

All mice underwent transthoracic echocardiography using a high-frequency ultrasound system with a 40-MHz linear transducer (Vevo 3100LT, Visual Sonics, Canada). In brief, mice were anaesthetized with isoflurane (in 100% O_2_ at 4% induction for 1 min and 1.5% for maintenance) and placed in supine position on a heated platform (37ºC). Heart rate was monitored along the recording period. After applying ultrasonic gel, the heart was visualized and a 2D M-mode video (4.5 seconds) was recorded from the parasternal short-axis view. To estimate the mitral valve inflow pattern, the transmitral LV outflow was also recorded from the apical four chamber view using the pulse wave (PW)-doppler imaging mode. All data was analyzed in Vevo 3100 Software, as previously described and according to the American Society of Echocardiography recommendations.^14^ The investigators performing and reading the echocardiograms were blinded to the treatment allocation.

**BM transplantation**

Donor animals (actin-eGFP) were euthanized and femurs were collected in cold PBS. Bone marrows were isolated from femurs and filtered through 40 μm cell strainers to obtain single cell suspensions. Depletion of mature hematopoietic cells, including T cells, B cells, monocytes/macrophages, granulocytes, erythrocytes and their committed precursors was achieved using a negative selection kit for Lineage Cell Depletion (Miltenyi). After washing, cell number and viability was assessed using an automated cell counter (Countess 3, ThermoFisher Scientific) and Trypan Blue solution (Merck, Germany). Cells were injected i.v. into Mx1^Cre^:c-Myb^fl/fl^ recipient mice (3-8x10^6^ cells per mouse) in a total volume of 100 μl saline. At the time of transplantation, recipient mice had previously been treated with poly(I:C) solution (Invivogen) at a dose of 10 μg g^-1^ body weight every other day for five times to induce BM depletion.^15^ Mice were maintained for 4 weeks after transplantation to establish efficient BM repopulation.

**Organ and tissue collection**

Mice were terminally anesthetized with ketamine (120mg/kg) and xylazine (16mg/kg) and blood was drawn via cardiac puncture and collected in 50mM EDTA tubes (Sigma-Aldrich). Plasma was isolated by centrifugation at 3000g for 10 min and stored at -80ºC until further use. Immediately after cardiac puncture, mice were transcardially perfused with 0.9% NaCl and the heart, both lungs, the right liver lobe, the spleen and both femurs were carefully excised and processed according to the specific endpoint.

**Cell isolation**

Spleen and bone marrow (from femur) were homogenized and filtered through 40 μm cell strainers to obtain single cell suspensions. Heart, lung and liver tissues were thoroughly minced in a digestive solution containing 60U/ml DNAse, 450U/ml Collagenase I, 125 U/ml Collagenase XI and 60 U/ml Hyaluronidase I-S in PBS and incubated for 30 min at 37°C on a shaker at 250 rpm. Afterwards, samples were filtered through a 40 μm cell strainer to obtain final single cell suspensions.

**Cell sorting**

Cell suspensions were obtained as described above and further purified by a density gradient centrifugation with Histopaque-107 at 300g, 4°C for 20 min. The mononuclear layer was isolated and cells were washed twice with PBS. Cell suspensions were stained with surface markers diluted in Brilliant Stain Buffer and sorted using a FACS Aria II cell sorter (BD Biosciences, Inc.) or a SH800S Cell Sorter (Sony Biotechnology). Propidium iodide (PI) was used as a cell viability marker.

**Flow cytometry**

The primary conjugated anti-mouse antibodies listed above were used for surface marker staining of different leukocytes subpopulations (see Key Resources Table). For high-dimensional flow cytometry of the heart, mice were injected i.v. with 3 μg CD45-e450 (see antibody list) 3 min before transcardiac perfusion, to exclude blood contamination. All samples were stained with Zombie NIR Fixable Viability Kit (1:1000) for 10 min at 4ºC and then with the specific surface markers diluted in Brilliant Stain Buffer, according to the manufacturer’s protocols, for 30 min at 4ºC. For the Click-iT™ EdU flow cytometry cell proliferation assay, prior to surface antibody staining, cell suspensions were fixed, permeabilized and the intracellular EdU was labelled by means of a click chemical reaction, following manufacturer´s instructions. All flow cytometric data was acquired using a Cytek Northern lights^TM^ flow cytometer (Cytek Biosciences, US) and analyzed using FlowJo software.

**Single cell RNA sequencing**

Three mouse single-cell RNA experiments were performed in this study: one on a pool of myeloid cells isolated from the heart, lung, liver, spleen and blood; and the other two on bone marrow myeloid cells, either from stroke and control mice, or from naïve recipients after BM transplantation. For the first experiment, mononuclear cell suspensions from peripheral organs were incubated with anti-CD16/CD32 antibody to block nonspecific binding and stained for CD45^+^CD11b^+^ myeloid cells (see antibodies list). Five unique cell hashtags antibodies were also used to label cells from each organ. All surface antibodies and hashtag antibodies were incubated for 30 min at 4ºC. Immediately prior to cell sorting, PI was added to all samples to label dead cells. Cells were sorted according to their surface markers (SH800S Cell Sorter, Sony Biotechnology). Sorted and organ-specific labeled cells from the heart, lung, liver, spleen and blood from the same animal were pooled together in the same collecting tube (cell ratio of 1:4:4:4:4, respectively). After sorting, all cells were centrifuged and cautiously resuspended to a final concentration of 1000 cells/μl. Cells were then transferred to the Next GEM chip according to manufacturer’s instructions. ScRNA-seq and cell hashing libraries were prepared using the 10x Chromium Single Cell 3′ Solution combined with feature barcoding technology for Cell surface protein (cell hashing), as per established protocols.^16^

For the second and third experiments, bone marrow cells suspensions were stained with a cocktail of antibodies against markers of the lymphoid lineage (CD3, CD4, CD8a, CD19 and Ter119) and mature neutrophils (Ly6G, see antibodies list). Immediately prior to cell sorting, PI or 7-AAD were also added to label dead cells. Cells were sorted according to the surface markers (SH800S Cell Sorter, Sony Biotechnology, or FACS Aria II Fusion cell sorter, BD Biosciences, Inc.) and the negative fraction of BM myeloid cells (depleted of lymphoid lineage cells and neutrophils) was collected. After sorting, cells were centrifuged and cautiously resuspended to a final concentration of 1000-1200 cells/μl. Cells were then further processed according to manufacturer’s (10X) instructions. The scRNA-seq libraries were prepared using the 10x Chromium Single Cell 3′ Solution.

In all three experiments, quality control of all cDNA samples was performed with a Bioanalyzer 2100 (Agilent Technologies) and libraries were quantified with the Qubit dsDNA HS kit. Gene expression libraries were sequenced on an Illumina NextSeq 2000 using 20,000 reads per cell. Cell-surface protein expression libraries were sequenced on an Illumina NextSeq 2000 aiming for 5,000 reads per cell.

**Single cell RNA-seq data processing and analysis**

Cell Ranger software was used to demultiplex samples, process raw data, align reads to the mouse mm10 reference genome (or a custom mouse mm10 reference genome with the eGFP marker gene incorporated) and summarize unique molecular identifier (UMI) counts. Filtered gene-barcode and hashing-barcode matrices that contained only barcodes with UMI counts that passed the threshold for cell detection were used for further analysis. Filtered UMI count matrices were processed using R and the R package Seurat. As quality control steps, the following cells were filtered out for further analysis: (1) doublets originating from two cells from different samples (one cell positive for two HTO); (2) cells with no HTO detected; (3) cells with a number of detected genes <500 or >6000; (4) cells with >7% of counts that belonged to mitochondrial genes. Hereafter, raw gene counts in high-quality singlets were log normalized and submitted to the identification of high variable genes by MeanVarPlot method. Data was scaled and regressed against the number of UMIs and mitochondrial RNA content per cell. Data was subjected to principal component analysis and unsupervised clustering by the Louvain clustering method. Cell clusters were visualized using Uniform Manifold Approximation and Projection (UMAP) representations. Clusters were manually annotated using the top upregulated genes for each cluster and through the UMAP visual inspection of the expression of key previously described markers. After initial cluster annotation, clusters of T and B cells (between 5-20% of total cells, depending on the experiment) were removed and all remaining myeloid cells were reanalyzed. Differentially expressed genes between conditions were calculated using the FindMarkers function. Volcano plots were generated using EnhancedVolcano in R (Bioconductor EnhancedVolcano v.1.6.0).^17^ Pathway enrichment analysis was performed using Ingenuity Pathway Analysis (Qiagen). Trajectory and pseudotime analysis were computed on the corresponding UMAP projections using Monocle 3.^18^

**Single Nuclei ATAC-seq**

Single nuclei suspensions were prepared as previously reported.^19^ In brief, BM cell suspensions were obtained as described for the single-cell sequencing experiments. Live, lymphoid lineage (CD3, CD4, CD8a, CD19 and Ter119) and neutrophil (Ly6G) negative BM cells were sorted using a FACS Aria II Fusion cell sorter (BD Biosciences, Inc.). Cells were centrifuged and incubated twice with Nuclei EZ Lysis Buffer on ice for 5 min each time. Lysed cells were washed and nuclei were incubated with 7AAD at 1μg/ml for microscopic inspection of integrity before sorting on a FACS Aria II Fusion (BD Biosciences). Then, nuclei were further processed according to manufacturer’s (10X) protocols at a concentration of 1000-5000 nuclei/µl. Single nuclei ATAC libraries were prepared using the 10x Chromium Next GEM Single Cell ATAC kit. Quality control of all cDNA samples was performed with a Bioanalyzer 2100 (Agilent Technologies). Libraries were sequenced on an Illumina NextSeq 1000, aiming for 25,000 reads per nuclei.

**Single nuclei ATAC-seq data processing and analysis**

Cell Ranger ATAC software (10x Genomics) was used to process raw single nuclei ATAC-seq data, align reads to the Mus musculus mm10 reference genome, generate the peak matrix with single-cell accessibility counts and the fragments file with unique fragments across all single cells. Filtered peak-barcode matrices that contained only barcodes with UMI counts that passed the threshold for cell detection were used for further analysis. Filtered matrices and fragment files were processed using R and the R packages Signac and Seurat. In brief, a chromatin accessibility matrix was created and as quality control steps, the following nuclei were filtered out for further analysis: (1) nuclei with a total detected fragments > 30,000 or < 5,000; (2) nuclei with the fraction of all fragments that fall within ATAC-seq peaks < 40%; (3) nuclei with a transcriptional start site (TSS) enrichment score lower than 2; and (4) nuclei with a ratio of mononucleosomal to nucleosome-free fragments higher than 4. Peaks were called using MACS2.^20^ Hereafter, UMAP based on latent semantic indexing (LSI) was generated to visualize data in the two-dimensional space and a gene activity matrix was created to quantify the activity of each gene in the genome by assessing its chromatin accessibility associated with each gene. Next, to help interpret data, the integration pipeline (canonical correlation analysis) from the Seurat package was used to perform cross-modality integration and label transfer on the single-cell ATAC dataset and single-cell mRNA sequencing dataset also generated from BM myeloid cells. Shared correlation patterns in the gene activity matrix and the single-cell mRNA sequencing were used to identify matched biological states and annotate predicted cluster labels for all nuclei from the ATAC object. Only those nuclei with a predicted score above the 0.5 cutoff were retained and considered for further analyses. Volcano plots showing differential accessible peaks between conditions were generated using EnhancedVolcano in R (Bioconductor EnhancedVolcano v.1.6.0).^17^ To identify differentially-active motifs between experimental conditions within specific cell types, we computed a per-cell motif activity score using chromVAR.^21^ For each gene, we used the ArchR package to identify set of peaks that regulate the gene by computing the correlation between gene expression and accessibility at nearby peaks. Peak-to-gene links were identified using default parameters, with k=100 and empirical p-value estimation. Positively correlated peak-to-gene links were defined with cutoffs r > 0.45 and FDR < 0.1. The corresponding peak-to-gene matrix was obtained by returning matrices from plotPeak2GeneHeatmap() function with k = 25, grouped by cluster identities. To further characterize these peak-to-gene links, the presence of putative transcription factor motifs was identified using the R package motifmatchr. Peaks were converted to GRanges and position weight matrices were obtained from JASPAR 2022 database.^22^

**Second harmonic generation microscopy imaging**

Mice were deeply anesthetized and euthanized as described above. Hearts were immediately extracted and immersed in 4% paraformaldehyde overnight. Afterwards, hearts were embedded in 4% agarose, coronally sectioned to 100 μm thick sections and kept at 4ºC as free-floating sections in PBS. Sections were then imaged using Zeiss LSM710 MT microscopy (Zeiss, Germany), acquiring second harmonic generation signals (445 nm) after excitation at a wavelength of 895 nm. SHG images were quantitatively analyzed with ImageJ software as previously described.^23,24^ In brief, the images underwent Fast Fourier Transformation (FFT) and then were submitted to an elliptic fit to obtain Aspect Ratio values as a measure of the anisotropy of the collagen fiber distribution.

**Mouse heart histological sections**

Mice were deeply anaesthetized and euthanized as described above. For conventional and immunofluorescence staining procedures, hearts were immediately extracted, submerged in 4% paraformaldehyde overnight at 4ºC, embedded in paraffin and coronally sectioned to 5 μm thick sections. For RNA-fluorescence in situ hybridization (RNA-FISH), hearts were excised, directly flash-frozen on dry ice and coronally cryosectioned to 5μm thick sections.

**Human heart histological sections**

Cardiac tissue specimens were obtained from adult subjects with ischemic stroke and controls. Specimens were immersed in 10% formalin upon collection to preserve tissue integrity, embedded in paraffin and sectioned at 3 μm thick sections.

**Picrosirius red staining**

Mouse and human heart paraffin sections were deparaffinized, stepwise in xylene, 100% ethanol, 70% ethanol and 50% ethanol, and then stained with Fast Green solution for 20 min. The sections were washed with 30% acetic acid, rinsed in tap water and submerged to Picrosirius red Solution for 60 min. Hereafter, sections were washed with 30% acetic acid and absolute alcohol, dried at room temperature (RT) and mounted with Eukitt® Quick-hardening mounting medium. Images were acquired at 20x magnification using Axio Imager 2 (Zeiss, Germany).

**Hematoxylin and eosin staining**

Mouse and human heart paraffin sections were processed and deparaffinized as previously described. Sections were incubated with Mayer's hematoxylin solution at RT for 5 min and washed under running tap water for 10 min. Afterwards, sections were incubated with 1% Eosin Y solution at RT for 3 min and rinsed with distilled water. Finally, sections were dehydrated stepwise with 70% ethanol, 80% ethanol, 90% ethanol and 100% ethanol, dried at RT and mounted with Eukitt® Quick-hardening mounting medium. Images were acquired at 100x magnification using Axio Imager 2 (Zeiss, Germany).

**Immunofluorescence staining**

Mouse heart paraffin sections were processed and deparaffinized as previously described. After deparaffinization, sections were incubated with blocking solution containing 2% goat serum, 1% BSA and 0.1% cold fish skin gelatin in PBS for 1 h at RT. Afterwards, sections were incubated with the primary antibodies against Collagen I (1:50), Collagen III (1:100) or Fibronectin (1:50) overnight at 4°C. Then, sections were washed in PBS and incubated with Alexa Flour 647 goat anti-rabbit secondary antibody (1:100) in the dark for 1 h at RT. After washing, sections were stained with DAPI and mounted with Fluoromount^TM^ Aqueous Mounting medium. Images were acquired in a confocal microscope at 40x magnification (LSM 880, LSM 980; Carl Zeiss, Germany)).

**Immunohistochemical staining**

Human heart paraffin sections were deparaffinized and submitted to antigen retrieval in sodium citrate buffer (pH 6) for 20 min at 90ºC. Then, sections were washed and immersed in 3% hydrogen peroxide in methanol for 15 min. Sections were then incubated with blocking solution containing 5% goat serum, 1% BSA and 0.1% cold fish skin gelatin in PBS for 1 h at RT, and afterwards with the primary antibodies against CD14 (1:100) or CCR2 (1:100) overnight at 4°C. After washing, sections were incubated with HRP-conjugated goat anti-rabbit secondary antibody (1:1000) in the dark for 1 h at RT. Finally, sections were washed, stained with Mayer’s hematoxylin for 5 min, dried and mounted with Fluoromount^TM^ Aqueous Mounting medium. Images were acquired at 20x and 40x magnifications using Axio Imager 2 (Zeiss, Germany).

**RNA-fluorescence in situ hybridization (RNA-FISH)**

The RNAscope Multiplex Fluorescent v2 was used on mouse flash-frozen and human paraffin sections, according to the manufacturer’s instructions. Mouse flash-frozen sections were first fixed with 4% PFA at 4°C for 30 min and dehydrated stepwise in 50% ethanol, 70% ethanol and 100% ethanol. Then, sections were incubated with protease IV at RT for 30 min and hybridized with probes specific to *Mmp9* or C*x3cr1* at 40°C for 2 h. Human paraffin sections were deparaffinized by incubating with Xylene for 10 min and then 100% ethanol for 5 min. Then, sections were incubated with Hydrogen peroxide for 10 min at RT and antigen retrieval was done by incubating sections with RNAscope® Retrieval Reagents at 95°C for 15 min. Sections were then incubated with protease III at RT for 30 min and hybridized with probes specific to MMP9 and CD14 at 40°C for 2 h. After hybridization, all sections (mouse and humans) were washed and incubated with a series of pre-amplifier and amplifier reagents, fluorophores and the HRP blocker at 40°C for 15–30 min each, according to manufacturers’ protocols. To detect the eGFP+ signal, specific mouse sections were further incubated with 0.5% Triton X-100 for 5 min, blocked with 4% BSA for 10 min at RT and incubated with 1:200 anti-GFP Nanobody conjugated to a fluorescent dye (GFP-booster) at 4°C overnight.  Finally, all sections were then stained with DAPI and mounted with Fluoromount^TM^ Aqueous Mounting medium. Images were acquired at 20x and 63x magnifications (Leica, Dmi8, Germany).

**MMP9 gelatin zymography**

Mice were deeply anaesthetized and euthanized as described above. Hearts were immediately extracted and flash-frozen on powdered dry ice. Frozen hearts were placed in microcentrifuge tubes containing RIPA lysis/extraction buffer already supplemented with protease/phosphatase inhibitor and 5mm steal beads, and placed on a tissue lyser (Qiagen, Germany) at 50HZ for 10 min. The total protein content of each sample was measured using the Pierce BCA protein assay kit. Same amounts of protein were loaded and fractioned on 10% Zymogram Plus (Gelatin) gels, according to the manufacturer’s instructions. Gels were then incubated in Zymogram Renaturing Buffer and in Zymogram Developing Buffer for 30 min each, both at RT with gentle agitation. Then, gels were incubated in Zymogram Developing Buffer overnight at 37°C and stained with the colloidal blue staining kit for 3 hours at RT. Gels were washed in water overnight and imaged with a gel scanner (Epson, Germany). Images were analyzed with ImageJ.

**Enzyme linked immunosorbent assay (ELISA)**

Total pro-MMP9 concentrations were measured in heart protein lysates using the pro-MMP9 Mouse ELISA kit, according to manufacturer’s (ThermoFisher) instructions. All samples were run in duplicates and all duplicates showed a coefficient of variation <15%. Pro-MMP9 concentrations are expressed per microgram of total protein content in the heart, measured using the Pierce BCA protein assay kit and following manufacturer’s instructions.

**qPCR**

Total RNA from FACS-sorted cells was extracted using Arcturus® PicoPure® RNA Isolation Kit, according to the manufacturer's instructions. Reverse transcription to cDNA was performed using High-Capacity Reverse Transcription Kit. qRT-PCR was performed with a standard SYBR-Green PCR kit protocol as previously described.^25^ Relative changes on *Mmp9* gene expression levels were normalized to *Ppia* gene expression levels by using the 2^-ΔΔCt^ method.^26^

**Splenocytes isolation and incubation with CVC**

Mice were deeply anaesthetized and euthanized as described above. Spleens were immediately removed and cell suspensions were prepared as previously described. Splenocytes were incubated for 2h at 37°C with CVC at different concentrations (0 μM, 5μM, 10 μM and 100 μM). Afterwards, cells were washed and stained with surface markers for flow cytometry as described above.

**Bulk mRNA sequencing of human heart samples**

Human heart paraffin sections were used to retrieve the PFA-cross-linked mRNA as previously described.^27^ In brief, PFA-fixed paraffin-embedded hearth tissue sections were scratched from each slide using a stereomicroscope (Olympus SZ51, Model# 1111260100), collected in cold PKD buffer supplemented with proteinase K solution and snap-frozen in liquid nitrogen until further use. To prepare the Smartseq2 libraries, samples were thawed at RT for 3 min and incubated at 56°C in a thermal cycler (lid temperature 66°C), for 4 h or until tissue was completely dissolved. Samples were then placed on ice and incubated at 56°C for 1 min with dT25 magnetic beads to reverse crosslinked samples. Samples were incubated at RT to allow mRNA hybridization and washed with 1x hybridization buffer (HB), containing 2x SSPE, 0.05% Tween-20 and 0.05% RNase Inhibitor. Samples were then washed with PBS with 0.1% RNase Inhibitor and incubated with RNase-free water for 2 min at 80°C to elute mRNA. Smartseq2 libraries were prepared from 1 ng mRNA, as previously described. Libraries were sequenced 2x60 reads base pairs paired-end on an Illumina NextSeq 1000 to a depth of 300,000-600,000 reads/sample.

**Bulk mRNA sequencing data processing and analysis**

FastQC was used to check the quality of fastq files. Low-quality reads and adapters were trimmed using Cutadapt using the following parameters: (1) reads <20 bp, and (2) quality cutoff of 20. The trimmed FASTQ files were mapped to the mm10 reference genome using STAR. The mapped reads (of lesion tissue sections) belonging to the same slide were merged using the merge BAM files tool in Galaxy version 4. To quantify the number of reads mapping to the exons of each gene, featureCounts program was used. Data was further analyzed in R. As quality control steps, the following samples were filtered out for further analysis: (1) Samples with a high percentage of mitochondrial genes (>20%); samples with a high percentage of ribosomal genes (>8%); (3) very highly-expressed genes with more than 20.000 counts; and (4) genes that are expressed in less than 5 samples with more than 1 count. Hereafter, raw gene counts were normalized to stabilize variance (regularized logarithm method) and differentially expressed genes were identified using the DESeq2 package in R.

**QUANTIFICATION AND STATISTICAL ANALYSIS**

Data were analyzed using GraphPad Prism version 9.0. All summary data are expressed as the mean ± standard deviation (s.d.), unless indicated otherwise. Normality was assessed in all datasets using the Shapiro-Wilk normality test. Normally-distributed data were analyzed using a two-way Student’s t test (for 2 groups) or ANOVA (for > 2 groups). Data with a no normal distribution were analyzed using the Mann-Whitney U test (for 2 groups) or Kruskal-Wallis test (H test, for > 2 groups). Multiple comparison adjusted p values were computed using Bonferroni correction or Dunn’s multiple comparison tests. A p value < 0.05 was considered statistically significant.

**SUPPLEMENTRY REFERENCES**

1.	Stuart, T., Butler, A., Hoffman, P., Hafemeister, C., Papalexi, E., Mauck, W.M., Hao, Y., Stoeckius, M., Smibert, P., and Satija, R. (2019). Comprehensive Integration of Single-Cell Data. Cell *177*, 1888-1902.e21. 10.1016/j.cell.2019.05.031.

2.	Stuart, T., Srivastava, A., Madad, S., Lareau, C.A., and Satija, R. (2021). Single-cell chromatin state analysis with Signac. Nat Methods *18*, 1333–1341. 10.1038/s41592-021-01282-5.

3.	Cao, J., Spielmann, M., Qiu, X., Huang, X., Ibrahim, D.M., Hill, A.J., Zhang, F., Mundlos, S., Christiansen, L., Steemers, F.J., et al. (2019). The single-cell transcriptional landscape of mammalian organogenesis. Nature *566*, 496–502. 10.1038/s41586-019-0969-x.

4.	Granja, J.M., Corces, M.R., Pierce, S.E., Bagdatli, S.T., Choudhry, H., Chang, H.Y., and Greenleaf, W.J. (2021). ArchR is a scalable software package for integrative single-cell chromatin accessibility analysis. Nat Genet *53*, 403–411. 10.1038/s41588-021-00790-6.

5.	Simon Andrews (2010). FastQC: a quality control tool for high throughput sequence data.

6.	Dobin, A., Davis, C.A., Schlesinger, F., Drenkow, J., Zaleski, C., Jha, S., Batut, P., Chaisson, M., and Gingeras, T.R. (2013). STAR: ultrafast universal RNA-seq aligner. Bioinformatics *29*, 15–21. 10.1093/bioinformatics/bts635.

7.	Marcel Martin (2011). Cutadapt Removes Adapter Sequences From High-Throughput Sequencing Reads.

8.	Liao, Y., Smyth, G.K., and Shi, W. (2014). featureCounts: an efficient general purpose program for assigning sequence reads to genomic features. Bioinformatics *30*, 923–930. 10.1093/bioinformatics/btt656.

9.	Love, M.I., Huber, W., and Anders, S. (2014). Moderated estimation of fold change and dispersion for RNA-seq data with DESeq2. Genome Biol *15*, 550. 10.1186/s13059-014-0550-8.

10.	Raudvere, U., Kolberg, L., Kuzmin, I., Arak, T., Adler, P., Peterson, H., and Vilo, J. (2019). g:Profiler: a web server for functional enrichment analysis and conversions of gene lists (2019 update). Nucleic Acids Res *47*, W191–W198. 10.1093/nar/gkz369.

11.	Percie du Sert, N., Hurst, V., Ahluwalia, A., Alam, S., Avey, M.T., Baker, M., Browne, W.J., Clark, A., Cuthill, I.C., Dirnagl, U., et al. (2020). The ARRIVE guidelines 2.0: Updated guidelines for reporting animal research. BMC Vet Res *16*, 242. 10.1186/s12917-020-02451-y.

12.	Llovera, G., Roth, S., Plesnila, N., Veltkamp, R., and Liesz, A. (2014). Modeling Stroke in Mice: Permanent Coagulation of the Distal Middle Cerebral Artery Video Link. J. Vis. Exp, 517293791–51729. 10.3791/51729.

13.	Kruger, A.J., Fuchs, B.C., Masia, R., Holmes, J.A., Salloum, S., Sojoodi, M., Ferreira, D.S., Rutledge, S.M., Caravan, P., Alatrakchi, N., et al. (2018). Prolonged cenicriviroc therapy reduces hepatic fibrosis despite steatohepatitis in a diet‐induced mouse model of nonalcoholic steatohepatitis. Hepatology Communications *2*, 529–545. 10.1002/hep4.1160.

14.	Lindsey, M.L., Kassiri, Z., Virag, J.A.I., De Castro Brás, L.E., and Scherrer-Crosbie, M. (2018). Guidelines for measuring cardiac physiology in mice. American Journal of Physiology - Heart and Circulatory Physiology *314*, H733–H752. 10.1152/ajpheart.00339.2017.

15.	Stremmel, C., Schuchert, R., Schneider, V., Weinberger, T., Thaler, R., Messerer, D., Helmer, S., Geissmann, F., Frampton, J., Massberg, S., et al. (2018). Inducible disruption of the c-myb gene allows allogeneic bone marrow transplantation without irradiation. Journal of Immunological Methods *457*, 66–72. 10.1016/j.jim.2018.03.016.

16.	Stoeckius, M., Zheng, S., Houck-Loomis, B., Hao, S., Yeung, B.Z., Mauck, W.M., Smibert, P., and Satija, R. (2018). Cell Hashing with barcoded antibodies enables multiplexing and doublet detection for single cell genomics. Genome Biology *19*, 1–12. 10.1186/s13059-018-1603-1.

17.	Blighe, K. & Sharmila Lewis, M. (2022). EnhancedVolcano: Publication-ready volcano plots with enhanced colouring and labeling. Bioconductor from within R. R version 1.8.0 (2020).

18.	Trapnell, C., Cacchiarelli, D., Grimsby, J., Pokharel, P., Li, S., Morse, M., Lennon, N.J., Livak, K.J., Mikkelsen, T.S., and Rinn, J.L. (2014). The dynamics and regulators of cell fate decisions are revealed by pseudotemporal ordering of single cells. Nat Biotechnol *32*, 381–386. 10.1038/nbt.2859.

19.	G Martelotto, L. (2019). ‘Frankenstein’ protocol for nuclei isolation from fresh and frozen tissue for snRNAseq v2 10.17504/protocols.io.3fkgjkw.

20.	Zhang, Y., Liu, T., Meyer, C.A., Eeckhoute, J., Johnson, D.S., Bernstein, B.E., Nusbaum, C., Myers, R.M., Brown, M., Li, W., et al. (2008). Model-based Analysis of ChIP-Seq (MACS). Genome Biol *9*, R137. 10.1186/gb-2008-9-9-r137.

21.	Schep, A.N., Wu, B., Buenrostro, J.D., and Greenleaf, W.J. (2017). chromVAR: inferring transcription-factor-associated accessibility from single-cell epigenomic data. Nat Methods *14*, 975–978. 10.1038/nmeth.4401.

22.	Castro-Mondragon, J.A., Riudavets-Puig, R., Rauluseviciute, I., Berhanu Lemma, R., Turchi, L., Blanc-Mathieu, R., Lucas, J., Boddie, P., Khan, A., Manosalva Pérez, N., et al. (2022). JASPAR 2022: the 9th release of the open-access database of transcription factor binding profiles. Nucleic Acids Research *50*, D165–D173. 10.1093/nar/gkab1113.

23.	Chiu, Y.-W. (2010). Second-harmonic generation imaging of collagen fibers in myocardium for atrial fibrillation diagnosis. J. Biomed. Opt *15*, 026002. 10.1117/1.3365943.

24.	Cicchi, R., Kapsokalyvas, D., De Giorgi, V., Maio, V., Van Wiechen, A., Massi, D., Lotti, T., and Pavone, F.S. (2009). Scoring of collagen organization in healthy and diseased human dermis by multiphoton microscopy. J. Biophoton. *3*, 34–43. 10.1002/jbio.200910062.

25.	Martinez-Fierro, M.L., Garza-Veloz, I., Castañeda-Lopez, M.E., Wasike, D., Castruita-De la Rosa, C., Rodriguez-Sanchez, I.P., Delgado-Enciso, I., and Flores-Mendoza, J. (2022). Evaluation of the Effect of the Fibroblast Growth Factor Type 2 (FGF-2) Administration on Placental Gene Expression in a Murine Model of Preeclampsia Induced by L-NAME. Int J Mol Sci *23*, 10129. 10.3390/ijms231710129.

26.	Livak, K.J., and Schmittgen, T.D. (2001). Analysis of relative gene expression data using real-time quantitative PCR and the 2(-Delta Delta C(T)) Method. Methods *25*, 402–408. 10.1006/meth.2001.1262.

27.	Ji, H., Besson-Girard, S., Androvic, P., Bulut, B., Liu, L., Wang, Y., and Gokce, O. (2023). High-Resolution RNA Sequencing from PFA-Fixed Microscopy Sections. In Neural Repair Methods in Molecular Biology., V. T. Karamyan and A. M. Stowe, eds. (Springer US), pp. 205–212. 10.1007/978-1-0716-2926-0_16.

28.	Kaya, T., Mattugini, N., Liu, L., Ji, H., Cantuti-Castelvetri, L., Wu, J., Schifferer, M., Groh, J., Martini, R., Besson-Girard, S., et al. (2022). CD8+ T cells induce interferon-responsive oligodendrocytes and microglia in white matter aging. Nat Neurosci *25*, 1446–1457. 10.1038/s41593-022-01183-6.
